# Glycolytic flux sustains human Th1 identity and effector function via STAT1 glycosylation

**DOI:** 10.1101/2025.02.06.636644

**Authors:** Ariful Haque Abir, Julia Benz, Benjamin Frey, Heiko Bruns, Udo S. Gaipl, Kilian Schober, Dimitrios Mougiakakos, Dirk Mielenz

**Author notes:** Correspondence: Prof. Dr. Dirk Mielenz Division of Molecular Immunology, Department of Internal Medicine 3, Friedrich-Alexander-Universität Erlangen-Nürnberg, Nikolaus-Fiebiger-Center, Glueckstr. 6, 91054 Erlangen, Germany.

## Abstract

Fate decisions of T helper (Th) cells are tightly linked to their metabolic states, but precise mechanistic links remain unknown, especially in humans. Using *in vitro* stimulation in combination with gene editing we studied how metabolic regulation shapes human Th1 cell identity and effector function. Differentiated Th1 cells displayed elevated STAT1 phosphorylation at Tyr701 and Ser727 as well as heightened T-bet and IFNγ expression, which were dampened by CRISPR/Cas9-mediated STAT1 deletion. Metabolic profiling revealed enhanced glycolytic activity in Th1 in comparison to Act.T cells, evidenced by increased extracellular acidification rate, ATP production via glycolysis, glucose uptake, lactate secretion and NADH abundance. SCENITH analysis demonstrated elevated glycolysis-dependent anabolic activity of Th1 cells. Inhibition of glycolysis reduced IFNγ production and STAT1 phosphorylation independent of JAK1/2 activity, STAT1 abundance or SHP-2 activity, implicating glycolysis directly in sustaining STAT1-mediated Th1 functionality. Mechanistically, O-Glycosylation, facilitated by O-Glycosyltransferase, emerged as pivotal in modulating STAT1 activity, as evident through immunoprecipitation and Western blot analysis. Pharmaceutical O-Glycosyltransferase inhibition prevented Th1 differentiation as well as STAT1 O-glycosylation. CRISPR/Cas9 mediated mutation of the O-glycosylation sites Ser499 and Thr510 sites diminished STAT1 Ser727 phosphorylation and IFNγ synthesis. Together, this study highlights glycolysis as key regulator of human Th1 cell identity and effector function, with STAT1 O-Glycosylation selectively maintaining Th1 effector capacity. This mechanism could be explored to safeguard Th1 cells in antiviral immunity and autoimmunity.

**Graphical abstract:** 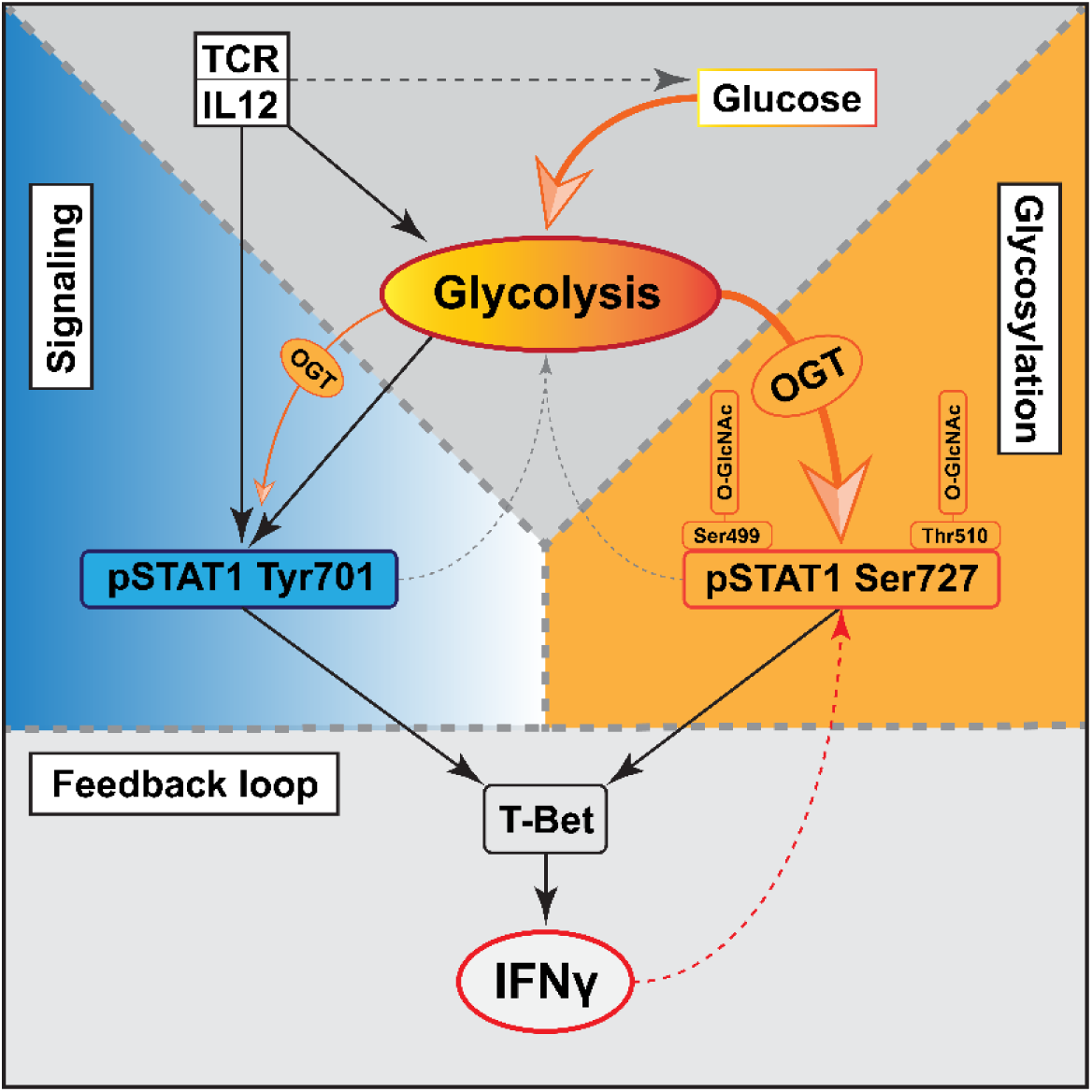

## Introduction

Naïve CD4^+^ T cells must receive signals from pMHC via CD3 and costimulation via CD28 for full activation (Kumar et al., 2018). The latter triggers anabolic phosphoinositol-3-kinase/Akt/mammalian target of rapamycin complex (PI3K/Akt/mTORC) 1 signaling, which is required for T cell growth and effector functions (Ho et al., 2009; Johnson et al., 2018). Seminal observations revealed that there are different Th effector cell types based on cytokine profiles (Mosmann et al., 1986). Th1 cells produce IFNγ, IL12 and TNFα while Th2 cells generate IL4, IL5, IL13; Th9 cells express IL9 and Th17 cells induce IL17 (Golubovskaya & Wu, 2016). IFNγ producing Th1 cells protect from intracellular bacteria, parasites, viruses and cancer. The downside is their involvement in autoimmunity (Annunziato & Romagnani, 2009; McNab et al., 2015). Thus, mechanisms controlling Th1 homeostasis and effector capacity, that is, IFNγ production, are of considerable interest. T-bet (TBX21) is the master transcription factor for Th1 cells and controls IFNγ production (Szabo et al., 2000, 2002). It reinforces its own expression, thereby, consolidating the Th1 phenotype (Afkarian et al., 2002; Lighvani et al., 2001). IL-12 has a pivotal upstream role in Th1 differentiation (Hsieh et al., 1993; Trinchieri et al., 1992) by inducing T-bet expression and Th1 polarization independent of IFNγ signaling. Hence, the autoregulation of T-bet becomes superfluous in the presence of IL-12 or IFNγ (Mullen et al., 2001). T-bet maintains Th1 identity by repressing the Th2 definining transcription factor GATA3 and vice versa (Usui et al., 2006). However, hybrid Th1/Th2 cells have been described in mice (Peine et al., 2013).

During Th1 differentiation, signal transducer and activator of transcription (STAT) 1 and Janus kinase (JAK) 1/2 take central stage (Shuai et al., 1993; Wilks, 1989; Wilks et al., 1991). STAT1 is essential for Th1 cell differentiation and the production of IFNγ. Phosphorylation of STAT1 via IL-12 mediated JAK2 activation at Tyr701 is the first step in the activation process (Barnholt et al., 2009; Seif et al., 2017); it enables the nuclear translocation of STAT1. Subsequent phosphorylation of STAT1 at Ser727 via autocrine IFNγ receptor signaling is essential for full transcriptional activation. JAK-mediated phosphorylation of STAT is a prerequisite to induce IFN-associated gene expression (Müller et al., 1993). In addition to phosphorylation at Tyr701 and Ser727, activation of STAT1 requires methylation of arginyl residues at the amino terminus which extends the half-life of STAT1 tyrosine phosphorylation (Subramaniam et al., 2001). Notably, Ser727 phosphorylation is independent of Tyr701, although both events are part of the IFNγ pathway (Zhu et al., 1997). Two splicing isoforms of STAT1 have been identified: STAT1 α (91 kDa) and β (84 kDa). Only STAT1α contains a full-length transcriptional activation domain with phosphorylation sites at Tyr701 and Ser727. In contrast, the STAT1β isoform has a truncated transcriptional activation domain and bears a single phosphorylation site at Tyr701 (Müller et al., 1993; Schindler et al., 1992).

CD4^+^ T cells mainly depend on OxPhos during their development (Kouidhi et al., 2017). However, activation via the PI3K/Akt/mTORC1 axis shifts the cells towards glycolysis (Kouidhi et al., 2017; Waickman & Powell, 2012). Of note, STAT1 transcriptionally activates glycolytic genes such as ENO1 and PDK3, further enhancing glycolysis, in human mesenchymal stem cells (MSC) (Jitschin et al., 2019). Glycolysis provides on the one hand rapid energy in the form of ATP and on the other hand supplies anaplerotic intermediates for redox balance and the hexosamine biosynthesis pathway. Major post-translational protein modifications supported by the hexosamine pathway are protein N- and O-glycosylation (Chiaradonna et al., 2018). Two enzymes, O-Glycosyltransferase (OGT) and O-Glycanase (OGA), add or remove O-glycosylation (Abramowitz & Hanover, 2018). O-Glycosylation is increased during human T cell activation, suggesting dominant OGT activity during this process (Lund et al., 2016).

Most studies focusing on Th1 cell metabolism are based on murine models. Given the importance of STAT1 in human Th1 cell function, and considering the metabolic implications of STAT1 signaling in MSC, we explored the intercalation of metabolism and STAT1 in human Th1 cells. We show that glycolytic flux drives human Th1 cell differentiation via STAT1 activity. OGT mediated glycosylation of STAT1 supports STAT1 phosphorylation at Ser727 in activated T cells, thereby stabilizing intracellular IFNγ production. Thus, glycolytic flux maintains OGT activity and STAT1 O-Glycosylation to sustain human Th1 cell effector. This finding could help to understand Th1 vulnerability in acute chronic viral infections or cancer, or assist Th1 targeted therapies in autoimmunity.

## Results

### STAT1 Activity and IFNγ production distinguish Th1 from activated human T cells

To understand how metabolism and STAT1 activity are connected in human Th1 cells, we established a human in vitro Th1 differentiation system, starting with magnetically purified naïve T cells out of human peripheral blood mononuclear cells (PBMC; Figure S1 and S2). To selectively distinguish T-bet driven Th1 cell metabolism from metabolic changes elicited by T cell activation via CD3/CD28, we included non-polarized activated T cells (Act.T) as controls. Consequently, Th1 polarization was identified by IFNγ staining (Figure 1a and 1b), with Th1 cultures containing ∼3.5 fold more IFNγ^+^ cells than Act.T cells. Moreover, Th1 cells produced more IFNγ but not IL-4 or IL-17 (Figure 1c, d, e). In line, T-bet was more abundant (Figure 1f, g). These data validate efficient human Th1 cell differentiation in our cultured cells and this setup was used throughout the study, unless mentioned otherwise.

**Figure 1:**
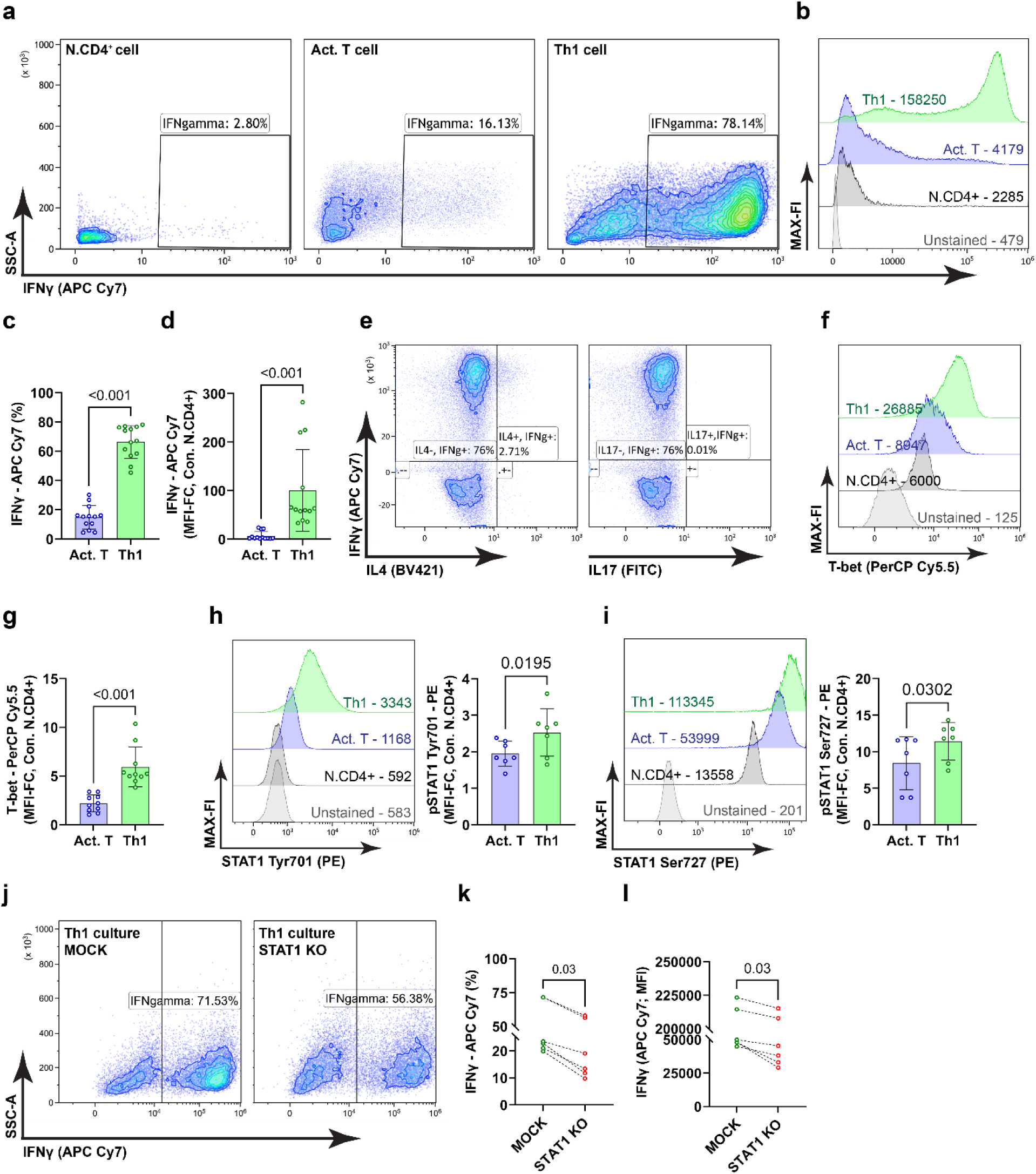
Analysis of IFNγ production and STAT1 Phosphorylation upon Th1 differentiation. Naïve CD4^+^CD45RA^+^ T cells (N.CD4^+^) were activated (Act.T cell) or Th1 polarized, intracellular staining was performed and cells were analyzed by flow cytometry. a. representative contour plots with IFNγ producing cells. b. representative histograms showing median fluorescence intensities (MFI) c. frequencies of IFNγ producing cells, mean ±SD, paired t-test, 13 donors, d. relative IFNγ abundance (Th1 vs. Act.T), mean ±SD, paired t-test, n=13, e. Th1 polarized cultures were intracellularly stained with anti-IL4 conjugated with Brilliant Violet 421 (BV421) and anti-IL17 conjugated with Fluorescein Isothiocyanate (FITC) and analyzed with flow cytometry. Representative contour plots are depicted. f. Th1 polarized cultures were intracellularly stained with anti-T-bet antibody conjugated with Peridinin chlorophyll protein-Cyanine5. 5 (PerCP Cy5.5) and analyzed by flow cytometry, representative histogram with MFI values. g. relative T-bet abundance (Act.T and Th1 cells va. N.CD4^+^), mean ±SD, paired t-test, n=10. h, i. STAT1 pTyr701 and pSer727 abundance, representative histograms and statistical analysis, mean ±SD, paired t-test, n=7. j,k,l. Th1 cultured cells were electroporated with STAT1-targeting Cas9-gRNA ribonucleoproteins (RNPs) or MOCK electroporated, and cells were further cultured for three days with polarizing medium. j, representative contour plots, k, l. frequencies of IFNγ producing cells and IFNγ abundance, paired t-test, mean ±SD, p=0.03, n=6.

Phospho-flow analysis of the IFNγ^+^ population in our culture corroborated the presence of high levels of STAT1 pTyr701 and pSer727 in Th1 vs. Act.T cells (Figure 1h and 1i). Therefore, efficient Th1 polarization is associated with an increase in the phosphorylation of STAT1 and the abundance of IFNγ and Tbet. To corroborate the function of STAT1 in our culture system, we induced a null mutation in the *stat1* gene in Th1 cultures by CRISPR/Cas9-mediated gene editing. Edited Th1 cultures contained lower frequencies of IFNγ positive cells and their IFNγ abundance was also lower (Figure 1 j-l). Together, these findings reinforce the notion of the involvement of STAT1 in human Th1 differentiation and effector function.

### Glycolytic Flux Contributes to Th1 Metabolism upon Differentiation

Murine Th1 cells shift towards aerobic glycolysis *in vitro* (Kouidhi et al., 2017; Palmer et al., 2015). To characterize metabolic changes during human Th1 cell differentiation, we performed extracellular flux (Seahorse) assays of naïve, Act.T and Th1 cell cultures. The extracellular acidification rate (ECAR) was upregulated in both Act.T and Th1 cells upon glucose addition, but ECAR was more strongly elevated in Th1 culture conditions (Figure 2a, b). The increase in glycolytic capacity and glycolytic reserve of Th1 cells indicated a higher rate of converting glucose into pyruvate or lactate (Figure 2c, d). ATP rate assays confirmed that Act.T, but even more so Th1 cells, produce their maximum ATP through glycolysis rather than by mitochondrial metabolism (Figure 2e).

**Figure 2:**
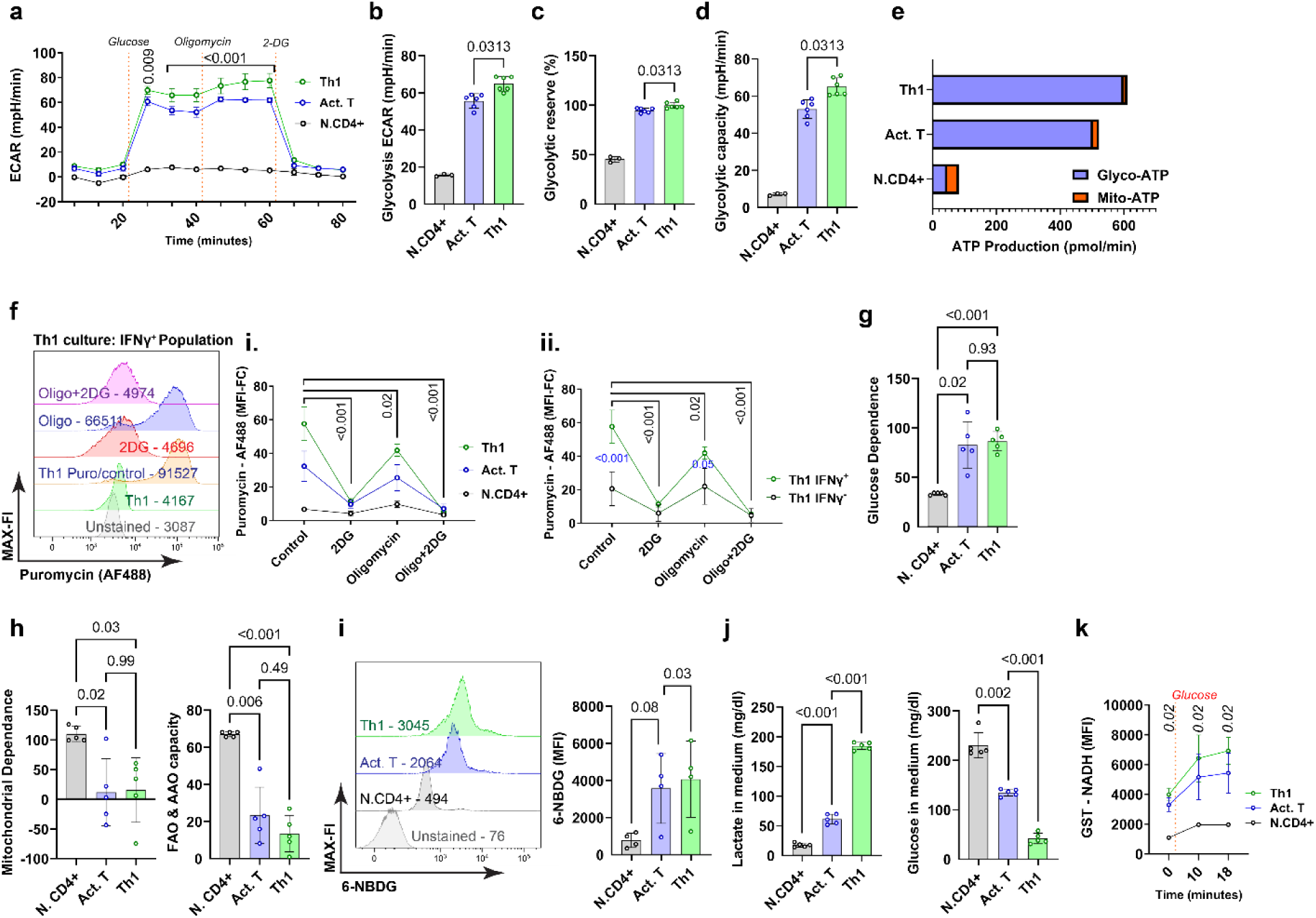
Metabolic characterization of human Th1 cells. a. N.CD4^+^, Act.T and Th1 cells were harvested after PMA and Ionomycin stimulation and Extracellular Flux Analysis was performed. The Extracellular Acidification Rate (ECAR) was measured using a Seahorse XF96 device using the Seahorse glycolysis stress test by serial injection of Glucose, Oligomicin and 2-Deoxy-D-glucose (2-DG). Statistical analysis was performed with two-way ANOVA repetitive measures, the black bar with p-value indicates the significance of the difference between Act.T and Th1 cells between 30 to 60 minutes, mean ±SD, n=6. b. ECAR (after glucose and before oligomycin injection), paired t-test, mean ±SD, n=6. c. Glycolytic reserve, paired t-test, d. Glycolytic capacity, paired t-test, e. Seahorse ATP rate assay, fi. representative histograms showing FI of anti-puromycin antibody conjugated with Alexa Fluor 488 (AF488) in different IFNγ^+^ populations in Th1 culture conditions; data are shown as mean±SD two-way ANOVA repetitive measures, the black bar with p-value indicates the significance of the difference of Th1 cell between controls and different inhibitors, n=5, fii: comparison of IFNγ+ and IFNγ-cells in Th1 cultures g, glucose dependence, one-way ANOVA with repetitive measures, h. mitochondrial dependence and fatty acid oxidation (FAO) and amino acid oxidation (AAO) were calculated using the SCENITH data, one-way ANOVA repetitive measures, n = 5. i. glucose uptake was measured by flow cytometry using 6-NBDG (6-(N-(7-Nitrobenz-2-oxa-1,3-diazol-4-yl)amino)-6-Deoxyglucose). MFI, value indicated in the representative histogram, of 6-NBDG was analyzed with one-way ANOVA repetitive measures, n=4, mean ±SD. j. medium was collected on the final day of culture after PMA and Ionomycin stimulation and glucose and lactate were quantified, one-way ANOVA repetitive measures, mean±SD, n=5, k. NADH fluorescence was measured after addition of glucose, two-way ANOVA repetitive measures, n=3, mean ±SD.

To support these data, flow cytometry-based single cell metabolic analysis was performed via SCENITH (Single Cell ENergetic metabolism by profiling Translation inHibition) in the same culture conditions (Argüello et al., 2020). SCENITH depicts the dependence of protein synthesis rate on the available ATP pool. The source of ATP is thereby quantified by measuring puromycin incorporation as a surrogate for protein synthesis, based on the assumption that protein translation is the most energy-demanding cellular process. Interestingly, Th1 cells exhibited a higher basal level of puromycin incorporation compared to Act.T (Figure 2f). This increased anabolic activity depended on glycolysis in both Act.T and Th1 cells (Figure 2g), much more so than on cellular respiration, which is blocked when oligomycin was used (Figure 2h). Of note, while protein translation in both Act.T and Th1 cell cultures depended on glycolysis, the reduction in puromycin incorporation by 2-deoxyglucose (2-DG) was stronger in Th1 cells (Figure 2fi), suggesting they are metabolically more active. Moreover, the IFNγ^+^ cells were more prone to 2-DG inhibition that the IFNγ^-^ cells in the same culture (Figure 2fii). Both Th1 and T.act cells depended less on fatty acid and amino acid oxidation than N.CD4^+^ cells (Figure 2h). Consistent with the Seahorse findings, these data support the hypothesis that anabolism of human Th1 cells is particularly fueled by glycolysis.

To substantiate these findings we performed a glucose uptake assay. After PMA/ionomycin restimulation, the cells were incubated in glucose-free medium with 6-NBDG (6-(*N*-(7-Nitrobenz-2-oxa-1,3-diazol-4-yl)amino)-6-Deoxyglucose), a fluorescent glucose analogon, for 30 min. Th1 cells showed only a slightly higher median fluorescence intensity (MFI) for 6-NBDG (Figure 2i). However, glucose and lactate abundance in the culture medium on day 5 of the culture period revealed that Th1 cultures contained much less glucose and more lactate, evidencing a strong glycolytic activity (Figure 2j). To monitor glucose driven NADH generation in real time we measured NADH by flow cytometry after adding glucose to starved cells (Abir et al., 2024) (Figure 2k), demonstrating that glycolytic flux is higher in Th1 than Act.T cells cells. Together, these six lines of evidence show that human Th1 cells exhibit a high biometabolic activitiy which mainly depends on glycolysis, with excess glucose turnover, suggesting a relationship between human Th1 differentiation and glycolysis.

### Glycolytic Flux Dictates Th1 Effector Function via STAT1 phosphorylation

To uncover the relationship between Th1 differentiation, that is, IFNγ production, and glycolysis, Th1 cells were treated short-term for 30 min with 2-DG and oligomycin, either separately or in combination, and IFNγ abundance was analyzed by flow cytometry. This analysis demonstrated that acute blockade of glycolytic flux with 2-DG notably reduced IFNγ production, without a discernible impact of oligomycin (Figure 3a). Furthermore, 2-DG diminished the frequency of IFNγ-producing cells (Figure 3b). These data indicate that glycolytic flux is essential to maintain both, Th1 identity as defined as the frequency of IFNγ-producing cells and effector capacity, defined as intracellular IFNγ amount.

**Figure 3:**
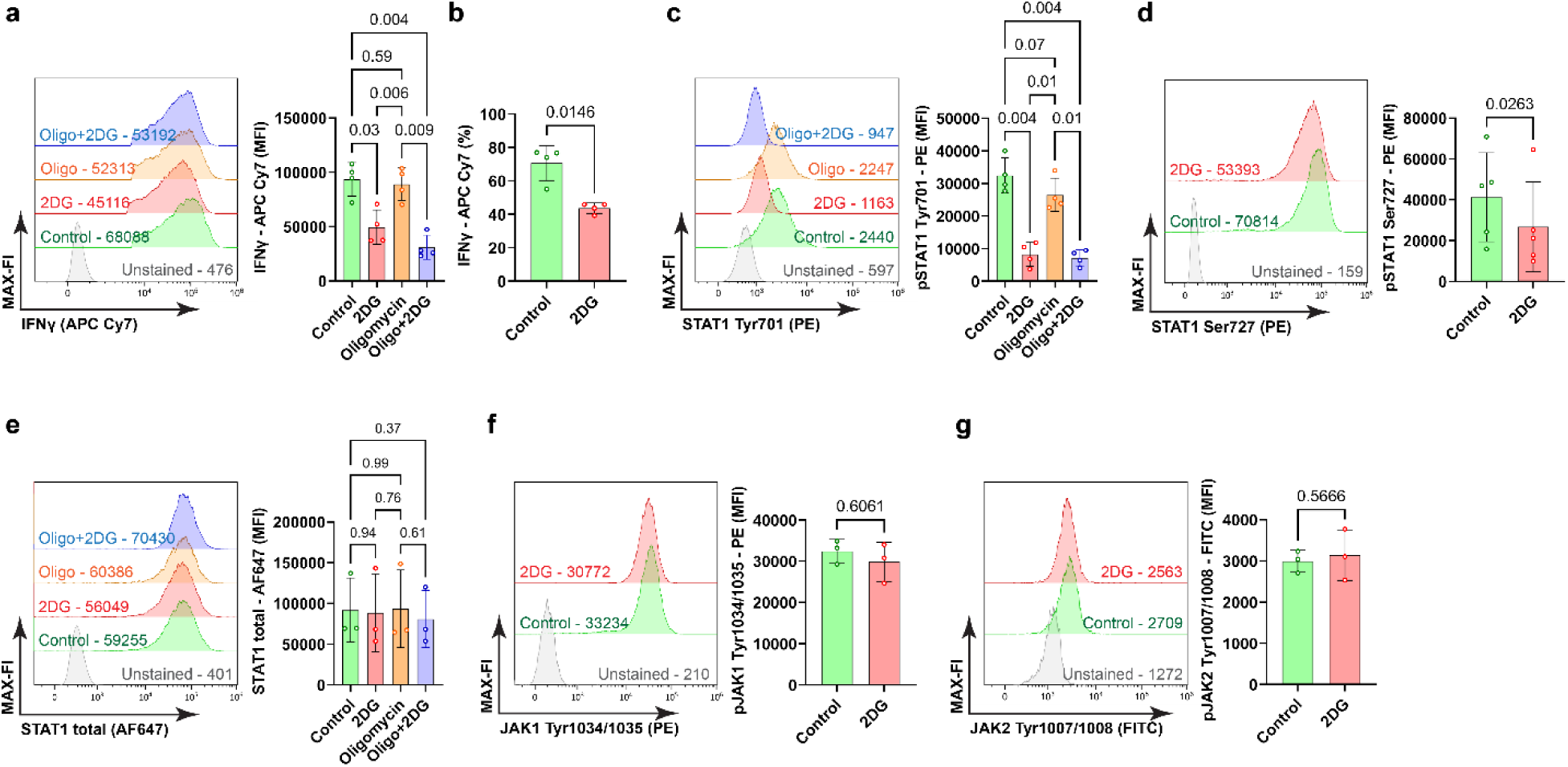
Th1 identity and effector capacity depend on glycolysis. Th1 cells were stimulated with PMA and Ionomycin for 30 min, treated with 2-DG and oligomycin either separately or in combination and were subjected to intracellular staining for flow cytometry analysis. a. Representative histogram shows MFI (representative value indicated) of anti-IFNγ conjugated with APC Cy7. One-way ANOVA with repetitive measures, n=4, mean±SD. b. frequency of IFNγ^+^ cells, a paired t-test, n=4, mean ±SD. c. representative histogram showing the MFI of anti-pSTAT1 Tyr701 conjugated with PE gated on IFNγ^+^ cells. One-way ANOVA with repetitive measures, n=4, mean±SD. d. representative histogram depicting MFI of anti-pSTAT1 Ser727 conjugated with PE gated on IFNγ^+^ cells, paired t-test, n=5, mean ±SD. e. representative histogram showing MFI of anti-STAT1 antibody conjugated with Alexa Fluor 647 (AF647) gated on IFNγ^+^ cells, one-way ANOVA with repetitive measures, n=3, mean±SD. f. representative histogram, MFI of anti-JAK1 Tyr1034/1035 conjugated with PE gated on IFNγ^+^ cells, paired t-test, n=3, mean ±SD. g. representative histogram, MFI, of anti-JAK2 Tyr1007/1008 conjugated with FITC gated on IFNγ+ cells, paired t-test, n=3, mean±SD.

The phosphorylation of STAT1 is a critical step in the signaling cascade that is necessary for the production of IFNγ in Th1 cells (Seif et al., 2017; Shuai et al., 1993). Accordingly, we investigated the impact of 2-DG on STAT1 activation. There was a clear reduction in STAT1 phosphorylation at both Tyr701 and Ser727 in response to glycolytic blockade (Figure 3c and 3d). To ascertain that these alterations were not influenced by upstream signaling or altered total STAT1 abundance, we measured total STAT1 levels and JAK1/2 phosphorylation at the activating residues Tyr1034/1035 and Tyr1007/1008, without significant changes (Figure 3e, f). These findings indicate that glycolysis plays a role in enhancing IFNγ production by modulating STAT1 phosphorylation downstream of JAK1/2 and independent of STAT1 abundance.

### 2-DG-induced STAT1 dephosphorylation partially depends on SHP2

Net phosphorylation is a balance between kinase and phosphatase activity. Src homology region 2 domain-containing phosphatase-2 (SHP2) can act as a dual-specific phosphatase in the case of STAT1, removing phosphorylation at both Tyr701 and Ser727, and 2-DG may intrinsically activate SHP2 (Chen et al., 2020, 2020; Wu et al., 2002). This could potentially contribute to the reduction of phospho-STAT1 in 2-DG-treated cells rather than the inhibition of glycolytic flux itself (Figure 4a). To test this possibility, Th1 cells were stimulated with PMA/ionomycin and co-treated with 2-DG as well as with two SHP2-specific inhibitors, TNO155 and RMC4550 (LaMarche et al., 2020; Pandey et al., 2019) (Figure 4a). Interestingly, SHP2 inhibition elevated the phosphorylation of STAT1 at Tyr701 and Ser727; however, this elevation of phosphorylation did not overcome completely the inhibition by 2-DG (Figure 4b and 4c). Hence, SHP2 inhibition only partially reverses the reduction of IFNγ-producing cells mediated by treatment with 2-DG. These findings indicate that besides SHP2 function, additional mechanisms downstream of glycolysis enforce STAT1 activity.

**Figure 4:**
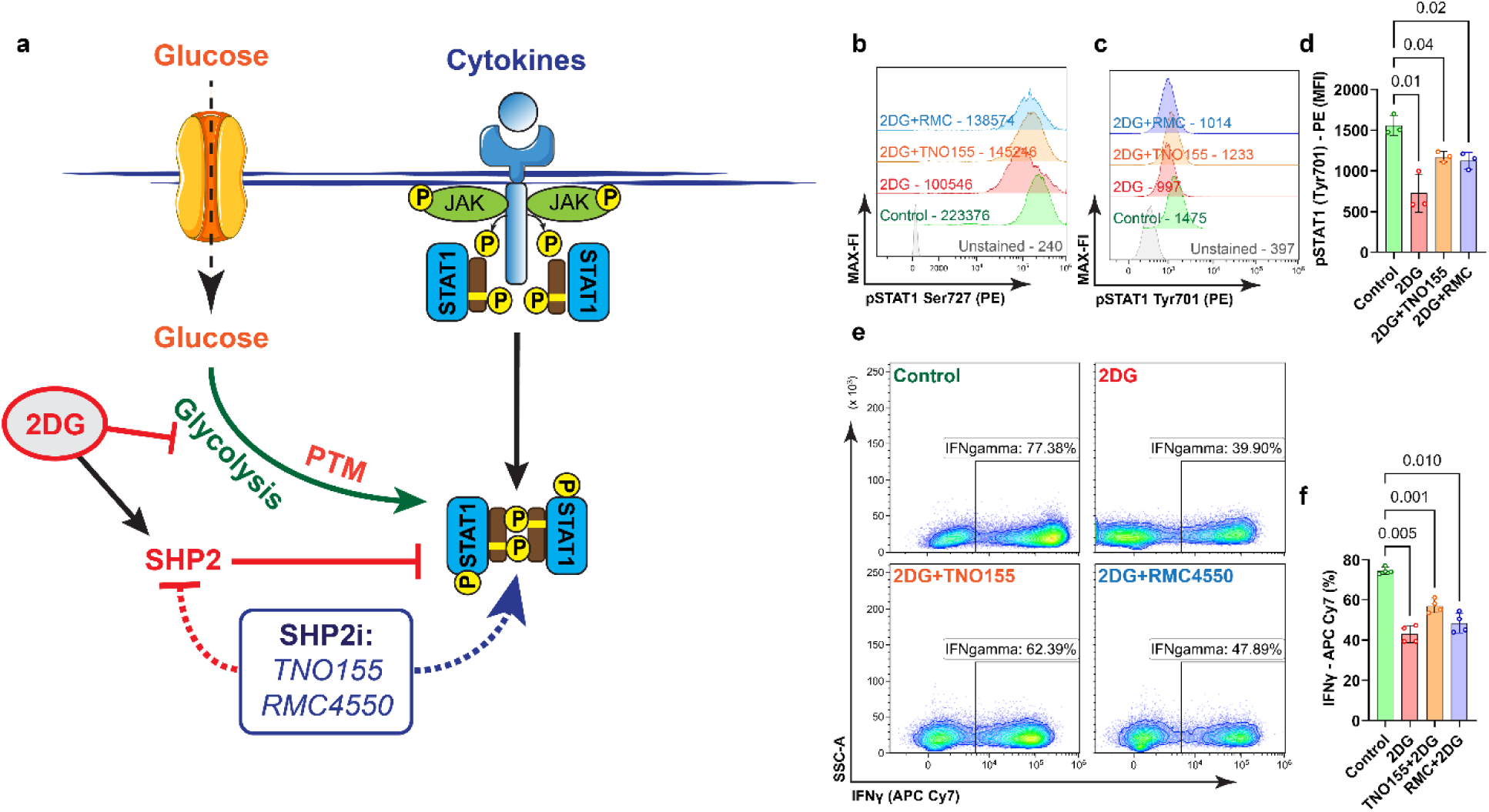
Glycolysis supports STAT1 phosphorylation and IFNγ abundance independent of SHP-2. a. Schematic depicting the experimental setup and read-out. SHP2 inhibitor (SHP2i), PTM, post-translational modification. PMA and Ionomycin-stimulated Th1 cells were further treated with 2-DG, TNO155, and RMC4550 in combination and intracellularly stained for flow cytometry analysis. b. representative histogram showing representative median FI (MFI) of anti-pSTAT1 Ser727 conjugated with PE gated on IFNγ^+^ cells. c. representative histogram with MFI of anti-pSTAT1 Tyr701 conjugated with PE from IFNγ^+^ cells, d. analysis of pSTAT1-PE MFI, one-way ANOVA with repetitive measures, n=3, mean±SD, e. frequencies of IFNγ^+^, representative contour plots. f. frequencies of IFNγ^+^ cells, one-way ANOVA repetitive measures, n=4, mean±SD.

### Glycolysis controls Th1 differentiation via STAT1 O-Glycosylation

To identify an additional mechanism by which glycolytic flux can influence STAT1 activity we considered the hexosamine biosynthesis pathway, which contributes the intermediate GlcNac to the O-Glycosylation pathway. First, we assessed the total O-Glycosylation status by intracellular staining with the anti-O-Glycosylation antibody RL2 (Jitschin et al., 2019). As described, T cell activation (Lund et al., 2016), but also Th1 differentiation increased global O-Glycosylation compared with naive T cell counterparts (Figure 5a), insinuating that O-glycosylation is mainly induced by activation and not by Th1 differentiation. Moreover, abundance of OGT was also augmented in both T.Act and Th1 cells (Figure 5b). However, this apparently does not exclude that O-glycosylation plays a role in Th1 identity, for instance, by targeting different substrates.

**Figure 5:**
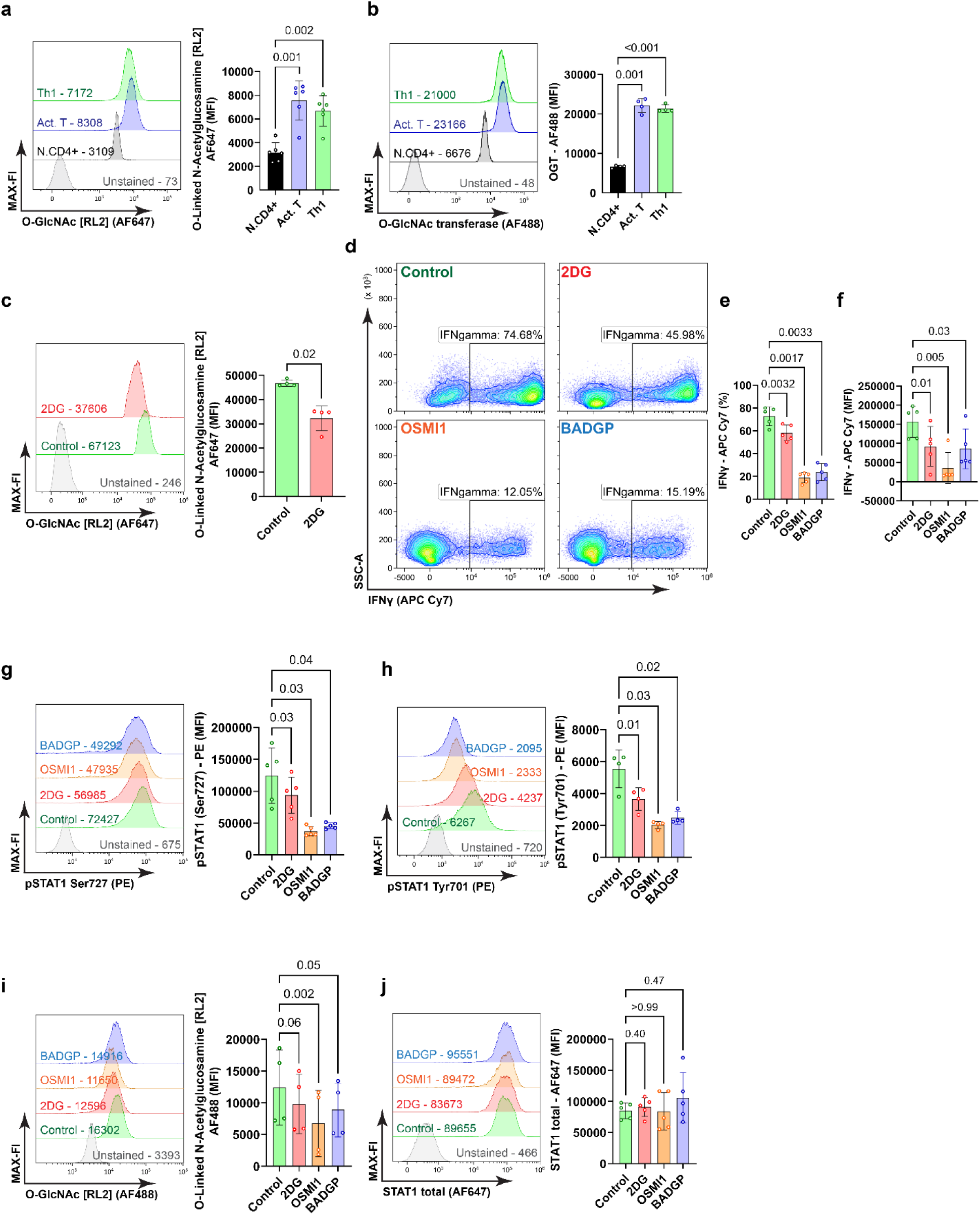
O-Glycosylation controls Th1 identity and effector capacity via STAT1 activity. a. Total O-glycosylation, representative histograms with representative median fluorescence intensity (MFI) of anti-O-Glycosylation antibody conjugated with Alexa Fluor 647 (AF647), one-way ANOVA with repetitive measures, n=6, mean±SD. b. Representative histograms with MFI of the anti-O-Glycosylation transferase antibody conjugated with Alexa Fluor 488 (AF488), one-way ANOVA with repetitive measures, n=4, mean±SD. c. Representative histograms with MFI of the anti-O-Glycosylation antibody conjugated with Alexa Fluor 488 (AF488) upon inhibition of glycolytic flux with 2DG. Paired t-test, n=4, mean±SD. d. Frequencies of IFNγ^+^ cells in control Th1 cells and 2-DG, OSMI1, and BADGP treated cells, representative contour plots. e. Frequencies of IFNγ^+^ cells, one-way ANOVA repetitive measures, n=5, mean±SD. f. MFI of IFNγ^+^, one-way ANOVA repetitive measures, n=5, mean±SD. g. Representative histograms with MFI of the anti-pSTAT1 Ser727 antibody conjugated with PE, one-way ANOVA with repetitive measures, n=5, mean±SD. h. Representative histograms with MFI of the anti-pSTAT1 Tyr701 antibody conjugated with PE, one-way ANOVA with repetitive measures, n=4, mean±SD. i. Representative histograms with MFI of the anti-O-Glycosylation antibody conjugated with AF647, one-way ANOVA with repetitive measures, n=4, mean±SD. j. A separate group of cells from the same setup was intracellularly stained with the anti-STAT1 antibody conjugated with Alexa Fluor 647 (AF647). Representative histograms with MFI, one-way ANOVA with repetitive measures, n=5, mean±SD.

To clarify the proportional relation between Th1 differentiation, O-glycosylation and glycolysis, Th1 cells were treated with 2-DG and subjected to the same analysis. 2-DG clearly reduced total O-Glycosylation in Th1 cells (Figure 5c). Pharmacological inhibition of OGT was then accomplished by adding two independent inhibitors, BADGP and OSMI1, in the last hour of PMA/ionomycin restimulation. This treatment revealed that OGT inhibition significantly decreased the number of IFNγ-producing cells and reduced the overall ability of IFNγ production (Figure 5d-f). Thus, similar to inhibition of Th1 cell differentiation with 2-DG, inhibition of protein O-Glycosylation reduces both, Th1 lineage commitment and effector function. In line, STAT1 phosphorylation on Tyr701 and Ser727 were significantly decreased after inhibition of OGT in the IFNγ^+^ population (Figure 5g, h), in line with recution of O-glycosylation (Figure 5i). However, there were no observable changes of total STAT1 presence between the experimental groups (Figure 5j). These data emphasize that O-Glycosylation mediated by OGT maintains Th1 cell identity and effector function by acting on STAT1 activity, indirectly or directly.

### Glycolysis and O-Glycosylation transferase mediate STAT1 O-glycosylation

The decrease of STAT1 phosphorylation in 2-DG-treated Th1 cells and in Th1 cells with OGT inhibition suggested an association between OGT activity and STAT1. To test whether O-Glycosylation occurs on STAT1 itself in Th1 cells, we immunoprecipitated (IP) STAT1 from Act.T and Th1 cells, respectively, and analyzed STAT1 O-Glycosylation by Western Blot. There was more O-glycosylated STAT1 in Th1 cells when compared to Act.T cells (Figure 6a, b). Next, we tested whether inhibition of glycolysis and OGT impairs STAT1 O-Glycosylation (Figure 6c, d). In fact, both inhibition of glycolysis with 2-DG, and OGT inhibition with BADGP and OSMI1, diminished STAT1 O-glycosylation. Together, these data support the hypothesis that OGT-mediated O-Glycosylation of STAT1 itself drives Th1 commitment and function.

**Figure 6:**
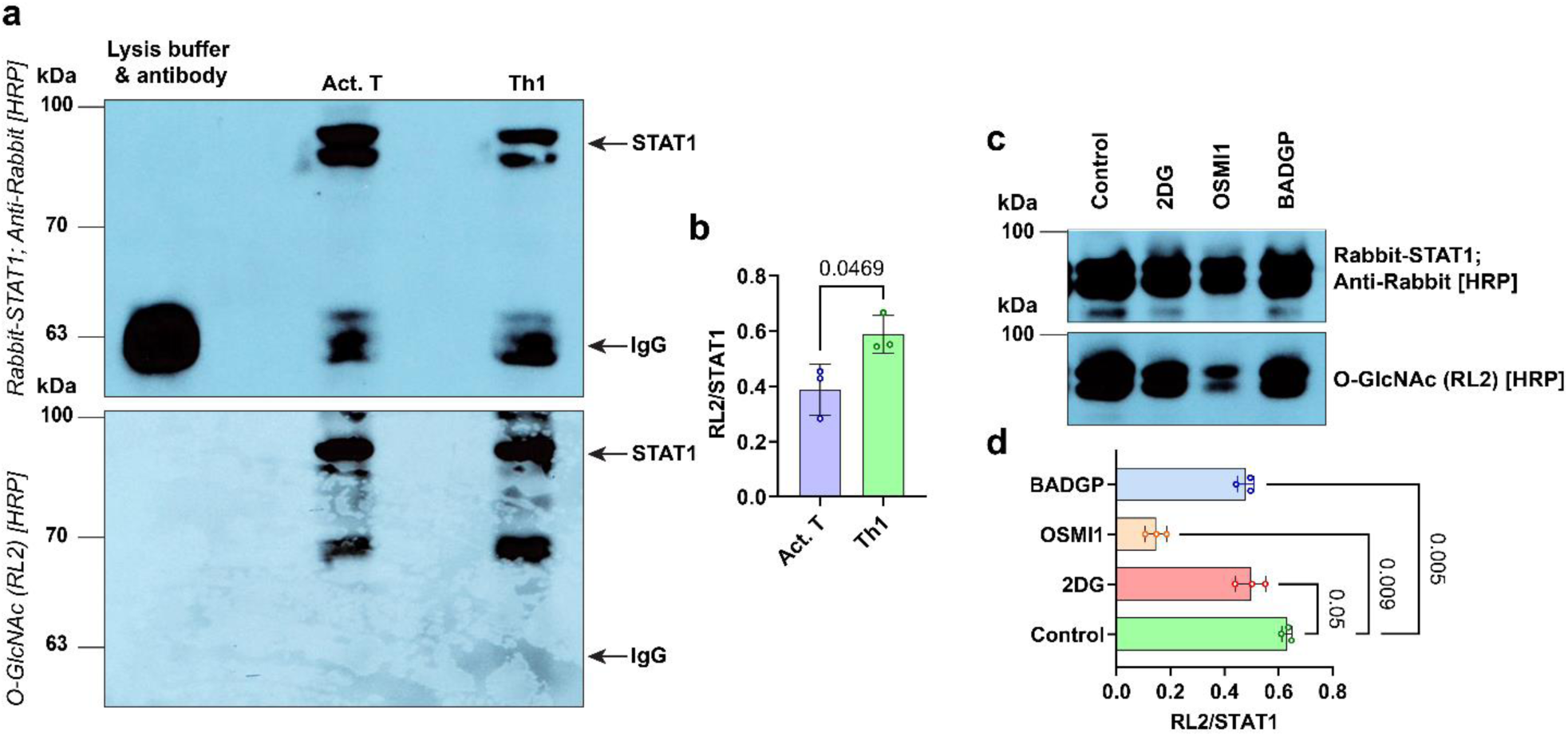
Analysis of STAT1 glycosylation (O-Glycosylation) a. Cells were harvested after PMA/Ionomycin stimulation, lysed, and STAT1 was immunoprecipitated (IP). Subsequently, samples were separated by 10% SDS-PAGE, followed by Western Blot. The membrane was stained with the horseradish peroxidase (HRP) conjugated anti-O-Glycosylation (RL2) antibody, washed and restained with a rabbit anti-STAT1 antibody, followed by anti-rabbit-HRP. The position of the STAT1 band and IgG are depicted. b. quantification of the intensity of the RL2/STAT1 signal ratio after scanning, paired t-test, n=3, mean±SD. c. Th1 cells were treated with 2-DG, OSMI1, and BADGP and were subjected to STAT1 IP, followed by western blot and membrane staining with STAT1 and RL2 detection. d. The intensity of the RL2/STAT1 signal ratio was calculated among the control (Th1), 2-DG, OSMI1, and BADGP treated groups, one-way ANOVA with repetitive measures, n=3, mean±SD. Molecular mass standards (kDa) on the left.

### STAT1 glycosylation is critical for lineage commitment

The previous experiment suggested that O-Glycosylation of STAT1 itself instructs Th1 lineage commitment as well as Th1 effector function. To test for an intrinsic STAT1 effect that mediates glycolysis- and OGT-driven Th1 effector identity and capacity, we analyzed the primary structure of STAT1. There are three predicted glycosylation sites, namely Thr489, Ser499, and Thr510 (https://glygen.org/protein/P42224#glycosylation). While Thr489 O-Glycosylation was identified once in Jurkat cells (Xu et al., 2022), Ser499 and Thr510 O-Glycosylation of STAT1 were identified in eight different human cell lines and primary cell types (Hahne et al., 2013; Jitschin et al., 2019; Y. Liu et al., 2020; Ramirez et al., 2021; Vang et al., 2024; Xie et al., 2021), most importantly also in activated human T cells (Lund et al., 2016); Woo et al., 2018). To validate the idea that intrinsic STAT1 glycosylation preserves Th1 lineage commitment and effector function, we replaced the two STAT1 glycosylation sites for which best evidence existed, Ser499 and Thr510, with Alanine via CRISPR/Cas9 mediated integration of a homology directed repair (HDR) template. A wild-type (unedited) STAT1 sequence was used as a control and a linked knock-in of GFP was used to identify the edited cells (Figure 7a; Table S1). T cells were activated in two ways: first, similar to all the previous experiments, IL12 was added directly after seeding to drive immediate Th1 cell differentiation. In order to establish Ser499Ala and Thr510Ala mutations before the onset of Th1 cell differentiation, Th1 cell differentiation was induced already on the 3^rd^ day after CRISPR editing. In this early differentiation setting, flow cytometry demonstrated reduced IFNγ abundance in STAT1 mutated (STAT1^mut^; GFP^+^) vs. unedited (GFP^-^) cells (Figure 7b) as well as less Th1 cells (Figure 7c). While the reduction of IFNγ abundance was mild in most cases, it was observed in 9 out of 10 analyses. The edited GFP^+^ STAT1^wt^ cells did not show decreased IFNγ abundance (Figure 7d) but also reduced Th1 cell differentiation (Figure 7e), most likely due to transient *stat1* gene inactivity during the editing process. Therefore, these data suggest that Ser499 and Thr510 may not be involved in Th1 cell differentiation but do regulate Th1 effector capacity. Similarly, IFNγ expression was lowered in edited STAT1^mut^ cells during late induction of differentiation (4 out of 6 experiments) while Th1 cell frequency was not affected (Figure S3). There was no effect on IFNγ production or Th1 cell frequency in edited STAT1^wt^ cells (Figure S3), indicating a potentially specific effect of STAT1^mut^ on IFNγ abundance in this setup. In line with the effect of STAT1^mut^ on IFNγ generation, analysis of STAT1 in the IFNγ^+^ population showed a significant reduction of pSer727 in all the cells with point mutations (GFP^+^) (Figure 7f; S3), irrespective of the differentiation protocol, while no change was observed at pTyr701 (Figure 7g). Moroever, edited STAT1^wt^ cells showed no reduction of pSer727 or pTyr701 (Figure 7h, i). Together, these data indicate that O-Glycosylation of STAT1 selectively stabilizes or fosters phosphorylation of STAT1 at Ser727, and, thereby, sustains Th1 cell effector function as defined by the intracellular amount of IFNγ.

**Figure 7:**
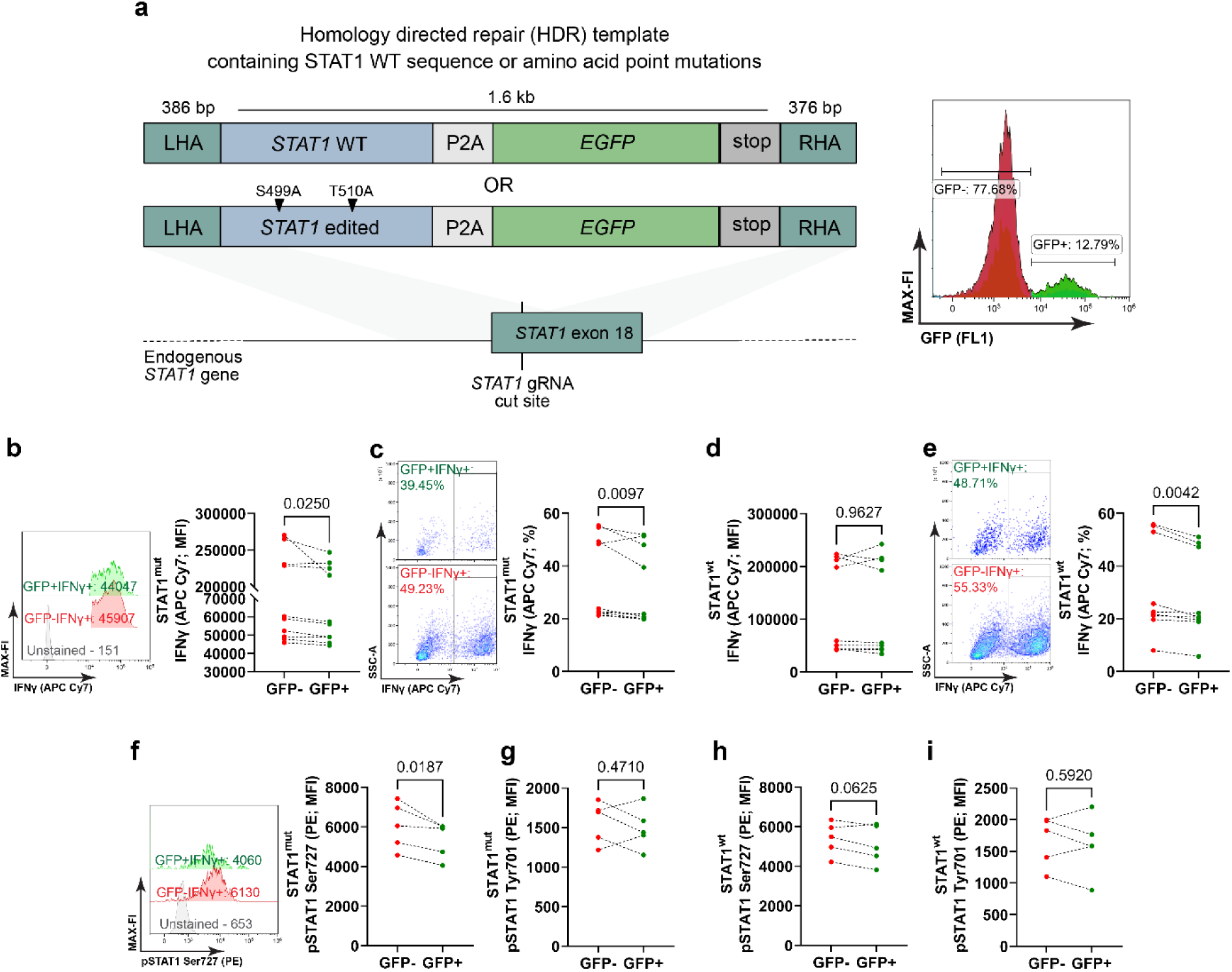
Analysis of Th1 effector function and STAT1 Phosphorylation upon mutation of the STAT1 glycosylation (O-Glycosylation) sites. a. Schematic of the *stat1* gene editing approach, LHA, left homology arm, RHA, right homology arm, representative histogram of electroporated cells. b. Representative histograms with MFI of the anti-IFNγ antibody conjugated with APC Cy7, GFP^+^ or GFP^-^ population of STAT1^mut^ cells (early differentiation), n=10. b. Representative contour plots of IFNγ producing cells, GFP^+^ or GFP^-^ populations of STAT1^mut^ cells (early differentiation), n=10. c. MFI of GFP⁺IFNγ⁺ and GFP⁻IFNγ⁺ cells from the STAT1^wt^ early differentiation group, n=9. d. Representative contour plots of IFNγ producing cells, GFP^+^ or GFP^-^ populations of STAT1^wt^ cells (early differentiation), n=9. e. Representative histograms with MFI of the anti-IFNγ antibody conjugated with APC Cy7, GFP^+^ or GFP^-^ populations of STAT1^mut^ cells (late differentiation), n=6. f. Frequencies of IFNγ producing cells gated from GFP^+^ or GFP^-^ populations of STAT1^mut^ cells (late differentiation), n=6. g. MFI of GFP⁺IFNγ⁺ and GFP⁻IFNγ⁺ cells from the STAT1^wt^ late differentiation group,n=6. h. Frequencies of IFNγ producing cells gated from GFP^+^ or GFP^-^ populations of STAT1^wt^ cells (late differentiation), n=6. i. Representative histograms with MFI of the anti-STAT1 Ser727 antibody conjugated with PE gated from GFP⁺IFNγ⁺ or GFP⁻IFNγ⁺ populations of STAT1^mut^ cells (early differentiation), n=5.

## Discussion

Mechanisms that intercalate metabolism and the signaling pathways which drive IFNγ production in primary human Th1 cells are scarcely reported. This study reveals that glycolytic flux and OGT activity both identify and determine Th1 lineage commitment as well as their their effector function. Downstream of glycolytic flux, OGT promotes STAT1 phosphorylation at Tyr710 and Ser727 as well as its O-Glycosylation. In accordance with reported O-Glycosylation of STAT1 at Ser499 and Thr510 in primary activated human T cells (Lund et al., 2016; Woo et al., 2018), we find that these two residues selectively fine-tune STAT1 controlled IFNγ abundance and Ser727 phosporylation. Thus, STAT1 O-Glycosylation, at least at Ser499 and Thr510, is not involved in initial IL-12 driven Th1 cell polarization. We therefore devise a model in which glycolysis and OGT-activity branch upstream of STAT1 O-Glycosylation. Together, these findings reveal glycolysis’ dual role in providing energy and substrates for post-translational modifications, ensuring Th1 lineage stability. Because even pharmacological short-term inhibition of glycolysis and OGT activity influenced IFNγ production and STAT1 phosphorylation, we propose that this mechanism equips Th1 cells to adapt dynamically to their environment; this could have important consequences for anti-tumor immunity. This notion is strongly supported by very recent data showing reduced IFNγ production in GLUT1-deficient human T cells or human T cells treated with a GLUT-1 inhibitor (Chen et al., 2023; Jong et al., 2024), yet we provide mechanistic insights downstream of glucose uptake.

In addition to the descriptive confirmation of the well established STAT1-IFNγ pathway in primary human T cells, we reinforced the relevance of this pathway in primary human T cells by Crispr/Cas9 mediated desctruction of the *stat1* gene. Compared to the mock control, frequencies of IFNγ producing cells and IFNγ abundance were reduced in the *stat1* edited cells. We provide several independent lines of evidence that dominant glycolytic metabolism distinguishes Act.T from Th1 cells. Functionally, inhibiting glycolysis for 30 min with 2-DG already reduced the frequency of IFNγ producing cells, IFNγ amount and STAT1 phosphorylation. Notably, 2-DG did neither affect signaling upstream of STAT1, such as JAK1/2 activation, nor total STAT1 protein abundance. However, glycolysis inhibition by 2-DG did reduce STAT1 phosphorylation by acting on SHP-2, which can dephosphorylate both Tyr710 and Ser727 (H. Chen et al., 2020, 2020; Wu et al., 2002). It is at present unclear whether glycolysis de-activates SHP-2 or whether the activating effect of 2-DG on SHP-2 is an off-target effect. Notwithstanding, SHP2 inhibition with two different inhibitors only partially restored STAT1 phosphorylation upon inhibition of glycolytic flux by 2-DG. These data therefore unravel additional regulatory mechanisms of IFNγ production besides STAT1 tyrosine phoshorylation. In fact, glycolysis increased OGT expression and O-Glycosylation in T.Act and Th1 cells and 2-DG reduced total O-Glycosylation. These findings align with previous work showing increased OGT activity in activated primary human T cells (Lund et al., 2016).

OGT activity was required to maintain STAT1 O-Glycosylation and phosphorylation as well as Th1 identiy and IFNγ production. Thus, OGT seems to be a major mediater of glycolytic pathways in T.Act and Th1 cells and it has reportedly many substrates in primary T cells and Jurkat cells (Lund et al., 2016; Woo et al., 2018; Xu et al., 2022). Of note, protein O-Glycosylation in contrast to protein N-glycosylation mainly targets cytoplasmic and nuclear proteins (Alfaro et al., 2012; Hart et al., 2007). O-Glycosylation can strongly enhance the half-life of proteins in the nucleus, more so in the cytoplasm (Xu et al., 2022), and it can influence nuclear-cytoplasmic shuttling (Hart et al., 2007). For instance, O-Glycosylation of nuclear factor κ B (NFκB) promotes its activation and nuclear translocation (Yang et al., 2008) while nuclear shuttling of nuclear factor of activated T cells (NFAT), another OGT substrate, is unaffected (Lund et al., 2016). On the other hand, lack of O-Glycosylation reduces STAT5’s tyrosine phosphorylation and transactivation potential (Freund et al., 2017). Consequently, O-Glycosylation of transcription factors exerts protein-specific roles. This apparent specificity is in line with the very specific functions we deduce for the function of STAT1 O-Glycosylation at Ser499 and Thr510, namely, maintenance of pSer727 and IFNγ amount. What could be further functional consequences of STAT1 O-Glycosylation? Glycosylation can protect transcription factors from proteasomal degradation or dephosphorylation (Han & Kudlow, 1997; D. Liu et al., 2002). An increased half-life would be in agreement with findings for other proteins (Xu et al., 2022). Yet, our data in T cells do not indicate major effects of O-Glycosylation on STAT1 protein half-life so far. It is possible that our experimental interventions were too short.

We extrapolate a proposed mechanism from CRISPR/Cas9 mediated gene editing. This was achieved by replacing Ser499 and Thr510 with Alanine residues. Importantly, off-target effects of the gene editing process were controlled for by Ser499/Ser and Thr510/Thr replacements by the HDR template. Because O-Glycosylation of STAT1 at Ser499 and Thr510 in primary human T cells is well established (Woo et al., 2018), we did not confirm the O-glycosylation of these particular residues. Still, we did corroborate that OGT targets STAT1 in Act.T cells, even more so in Th1 cells. It was suprising to see that Ser499 and Thr510 editing merely affects Ser727 phosphorylation and not Tyr510 phosphorylation. This specificity represents an important signaling loop for Th1 identity controlled by glycolytic flux. There may remain a putative role of Thr489 O-Glycosylation. However, in stark contrast to Ser499 and Thr510, O-Glycosylation of Thr489 was only found once at all, in Jurkat cells (Xu et al., 2022) and never in primary human T cells (Lund et al., 2016; Woo et al., 2018) or other cell types (https://glygen.org/protein/P42224#glycosylation).

The mechanism that we describe here may imprint inflammatory priming of Act.T cells by IL-12 and, consequently, present a potential therapeutic target that may help to limit inflammation or booster anti-viral or anti-tumour effector function. This suggestion aligns with the inflammatory priming in MSC in which a similar mechanism was described, albeit, in MSC, STAT1 glycosylation elicits an immunoregulatory phenotype (Jitschin et al., 2019). Because the mechanism of glycolysis and OGT-driven STAT1 glycosylation appears to be shared in human MSC and Th1 cells, this signaling circuit might be a general and conserved pathway, yet with cell type specific outcomes. Future experiments should aim at identifying O-Glycosylation dependent STAT1 target genes in different cell types. Our study underscores the previous finding that STAT1 O-Glycosylation integrates nutrient sensing with immunoregulatory processes in MSCs, forming a mechanism for controlling inflammation (Jitschin et al., 2019). IL-12 is a crucial for Th1 cell polarization and its abundance as well as polymorphisms of IL-12 and its receptor are linked to various inflammatory disease (Tan et al., 2009). Along this line, overnutrition, such as hyperglycemia, can support inflammation as well. It is tempting to speculate that continuous availability of gluocse creates a self-sustaining pro-inflammatory loop via enhanced glycolysis and O-Glycosylation in Th1 cells. Of note, glucose and other nutrients also induce the mTOR pathway. Interestingly, there is crosstalk between mTOR and STAT signaling (Jitschin et al., 2019; Kristof et al., 2003; Kroczynska et al., 2009; Ramana et al., 2002). The specific mode of interaction between mTOR and STAT1 signaling pathways requires further investigation but STAT1 O-Glycosylation might contribute to the interplay, albeit this is speculative at the moment. In conclusion, our observations demonstrate a mechanistic connection between glycolysis, STAT1 phosphorylation, and its O-Glycosylation during human Th1 differentiation and effector capacity.

## Author contribution

Data presented in this study were part of the doctoral thesis of Ariful Haque Abir. AHA. designed, performed and analyzed the experiments and wrote the paper; JB designed and performed experiments; HB, UG and BJ provided human PBMCs and edited the manuscript; KS designed experiments and edited the paper; DMo designed experiments and edited the paper; DMi designed and analyzed experiments and wrote the paper.

## Declaration of conflict of interest

The authors declare no conflict of interest

## Supporting information

Abir_2025_Supplement

## Acknowledgement

We acknowledge all voluntary healthy donors. We thank Profs. Hyun-Dong Chan and Christina Zielinski for helpful discussions.

## Funding

The study was funded by the Deutsche Forschungsgemeinschaft (DFG) through the Research Training Grant (RTG) 2599, P02 (to D.Mi.) and P08 (to D.Mo.), FOR2886 “PANDORA”, P03, to D.M., a Start-up grant through the RTG2599 (to A.H.A.), FOR5560, P08, to D.M.

## Methods (continued in the Supplement)

### Human Peripheral blood mononuclear cells

Peripheral blood mononuclear cells (PBMC) were freshly isolated via density gradient centrifugation from leukoreduction system chambers (LRSC) of healthy human donors. The permission to use this LRSC from Department of Transfusion Medicine and Haemostaseology at Universitätsklinikum Erlangen was given by the ethics committee of the Friedrich-Alexander-Universität Erlangen-Nürnberg (ethical approval no. 48_19 B and 21-400-Bp). The blood was diluted 1:1 with PBS and added to a tube containing 15 ml of Pancoll human (density: 1.077 g/ml). After centrifugation, the peripheral blood mononuclear cells (PBMCs) were collected from the interface of the Pancoll/erythrocyte mixture and serum. The PBMCs were cryopreserved in the vapor phase of liquid nitrogen. The samples were subsequently thawed and utilized for further experiments.

### Naïve CD4^+^ T cell Isolation

Naive CD4^+^CD45RA^+^ T cells were isolated from thawed PBMC with the Human Naive CD4^+^ T Cell Isolation Kit II (from Miltenyi Biotec, at. No.: 130-094-131) via magnetic cell sorting (MACS). In brief, PBMC were washed with the MACS buffer (phosphate-buffered saline or PBS, pH 7.2, 0.5% human serum albumin (HAS), and 2 mM ethylenediaminetetraacetic acid or EDTA).10 μl of MACS Naive CD4^+^ T Cell Biotin-Antibody Cocktail II, human, were added per 10⁷ cells and incubated for 5 minutes at 4°C. Subsequently, 20 μL of Naive CD4^+^ T Cell MicroBead Cocktail II were added per 10⁷ total cells, followed by the addition of some MACS buffer. Following a 10-minute incubation period, the cells were passed through a MACS LS column (Miltenyi Biotec; Cat. No.: 130-042-401) and placed into a MACS magnet system (QuadroMACS™ Separator; Miltenyi Biotec; Cat. No.: 130-091-051). Both the column and the magnet system were pre-cooled to 4°C. The isolated untouched cells were collected into a Falcon tube for subsequent culture. A small sample of cells was analyzed to determine the purity of the CD4^+^CD45RA^+^ population via flow cytometry. The accepted purity for the Th1 culture system was between 96% and 98%.

### Cell culture

The culture was set up in a 96-well plate coated with Ultra-LEAF™ anti-CD3 (3 μg/ml) (BioLegend; Cat. No.: 300438) and anti-CD28 (5 μg/ml) (BioLegend; Cat. No.: 302934) antibodies in 50 μl PBS per well. Following a two-hour incubation period at 37°C, the plates were aspirated, and cells seeded at a density of 50,000 cells/well in a volume of 200μl. Standard R10 medium was prepared and used for the culture by combining 500ml of RPMI1640 (Gibco™, Thermo Fisher Scientific; Cat. No.: 11875093) with 10% heat-inactivated fetal calf serum (FCS), 2mM L-glutamine, 50 U/ml penicillin G, 50 μg/ml streptomycin, and 50 μM β-mercaptoethanol. Activated and Th1 differentiation cultures were supplemented with 8 ng/ml IL-2 (Miltenyi Biotec; Cat. No.:130-097-743) and 10 μg/ml anti-IL-4 (BioLegend; Cat. No.:500802). Furthermore, the Th1 cultures were supplemented with 8 ng/ml IL12, to promote differentiation. Following a 48-hour incubation period at 37°C with 5% CO₂, the Th1 culture was transferred to a fresh plate. Following an additional 72-hour incubation period, the cells were restimulated with 10ng/ml PMA (Abcam; Cat. No.:ab120297) and 500ng/ml ionomycin (Sigma-Aldrich; Cat. No.: I3909) with 1µl/ml GolgiPlug (BD Biosciences; Cat. No.: 555029) for four hours before flow cytometry analysis. Unless otherwise indicated, this culture system was used for all experiments.

### Flow cytometry

#### Intracellular staining

Approximately 2×10^5^ cells were collected and rinsed with cold PBS. Subsequently, the cells were stained with Zombie Aqua (1:1000; Fixable Viability Kit, BioLegend; Cat. No.: 423101) for 15 minutes at room temperature in the dark. Then, the cells were washed twice with PBS and 400 μl of CytoFix buffer (preheated at 37°C; BD Cytofix™ Fixation Buffer; BD; Cat. No.: 554655) was added to each sample, followed by a 10-minute incubation at 37°C. Following a wash with cold PBS, the cells were permeabilized using PhosFlow buffer (precooled at -20°C; BD Phosflow™ Perm Buffer III; BD; Cat. No.: 558050) and incubated at 4°C for 30 minutes. After that the cells were washed twice with FACS buffer (PBS + 0.25% BSA) and incubated for 10 minutes with BD Fc Block™ (BD, Cat. No.: 564219, dilution: 1:100). The cells were then treated with the corresponding antibodies, as detailed in Table 1. Following a 30-minute incubation period with the antibodies, the cells were washed twice. The flow cytometry analysis was conducted using a Beckman Coulter Gallios Flow Cytometry system, and the resulting data were analyzed using the Beckman Coulter Kaluza Analysis Software.

### Flow cytometry antibody panels

#### Panel 1: Analysis of cell purity after MACS isolation

| Antibody | Concentration |
| --- | --- |
| APC anti-human CD4 Antibody | 1:100 |
| FITC anti-human CD45RA Antibody | 1:100 |

#### Panel 2: Analysis of Th1 differentiation

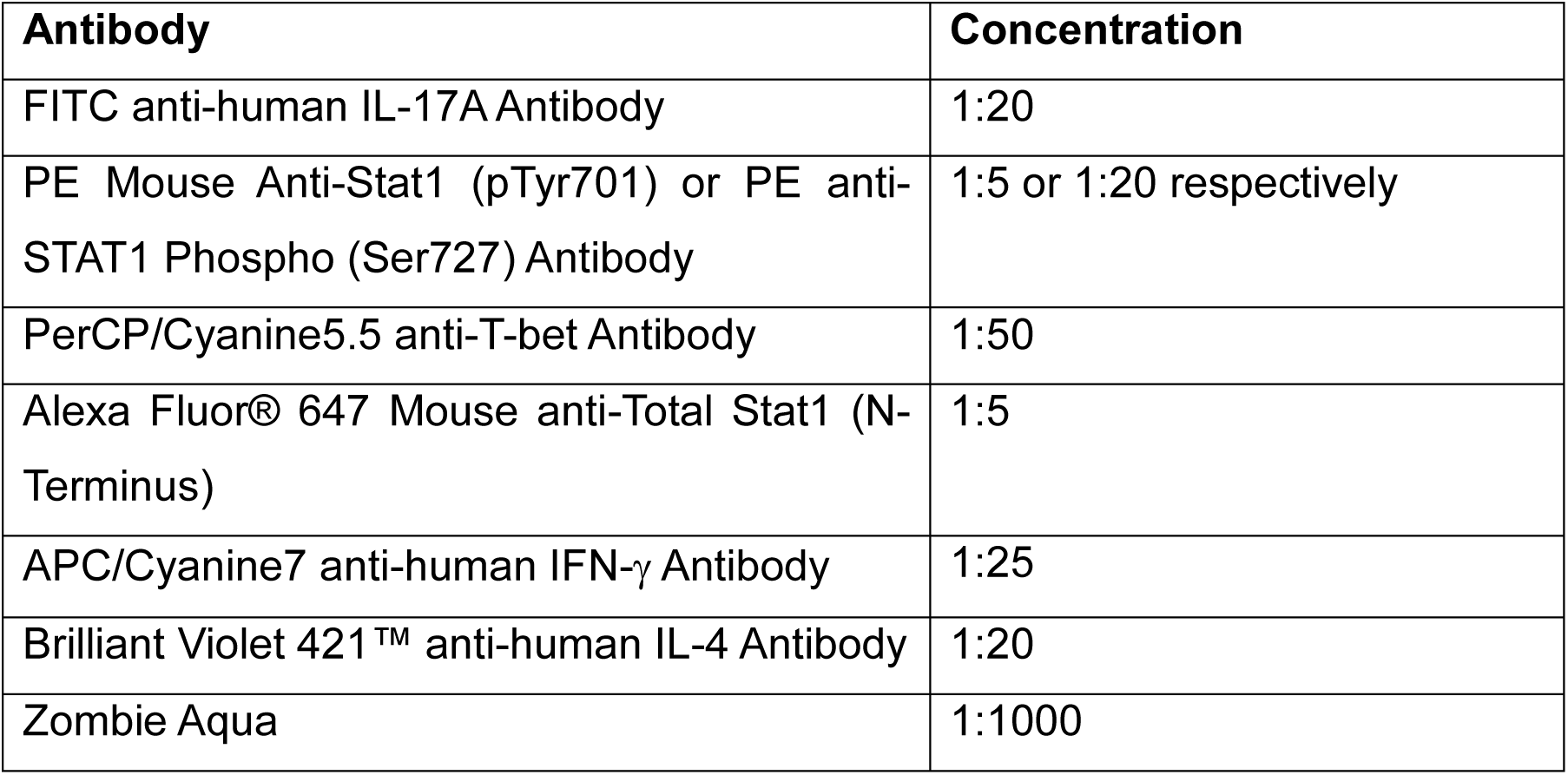

| Antibody | Concentration |
| --- | --- |
| FITC anti-human IL-17A Antibody | 1:20 |
| PE Mouse Anti-Stat1 (pTyr701) or PE anti-STAT1 Phospho (Ser727) Antibody | 1:5 or 1:20 respectively |
| PerCP/Cyanine5.5 anti-T-bet Antibody | 1:50 |
| Alexa Fluor® 647 Mouse anti-Total Stat1 (N-Terminus) | 1:5 |
| APC/Cyanine7 anti-human IFN-γ Antibody | 1:25 |
| Brilliant Violet 421™ anti-human IL-4 Antibody | 1:20 |
| Zombie Aqua | 1:1000 |

#### Panel 3: Analysis of STAT1 upstream signaling analysis

| Antibody | Concentration |
| --- | --- |
| FITC Phospho-Jak2 (Tyr1007, Tyr1008) Recombinant Rabbit Monoclonal Antibody | 1:20 |
| PE Phospho-Jak1 (Tyr1022, Tyr1023) Recombinant Rabbit Monoclonal Antibody | 1:20 |
| PerCP/Cyanine5.5 anti-T-bet Antibody | 1:50 |
| Alexa Fluor® 647 Mouse anti-Total Stat1 (N-Terminus) | 1:5 |
| APC/Cyanine7 anti-human IFN-γ Antibody | 1:25 |
| Zombie Aqua | 1:1000 |

#### Panel 4: Analysis of O-Glycosylation

| Antibody | Concentration |
| --- | --- |
| Alexa Fluor 488 Anti-O-Glycosylation Monoclonal Antibody [RL2] or<br>Alexa Fluor 647 Anti-O-Glycosylation [RL2] * | 1:20 or 1:100 respectively |
| FITC OGT1 Polyclonal Antibody * | 1:100 |
| PE Mouse Anti-Stat1 (pTyr701) or PE anti-STAT1 Phospho (Ser727) Antibody | 1:5 or 1:20 respectively |
| PerCP/Cyanine5.5 anti-T-bet Antibody | 1:50 |
| APC/Cyanine7 anti-human IFN-γ Antibody | 1:25 |
| Zombie Aqua | 1:1000 |

\* only used with Alexa Fluor 647 Anti-O-Glycosylation [RL2], otherwise in a separate set.

### SCENITH: Single Cell ENergetIc metabolism by profiling Translation inhibition

The SCENITH experiment was conducted as previously described (Argüello et al., 2020), with minor modifications. In summary, approximately 1 x 10⁶ naïve CD4⁺, activated T, and Th1 cells were restimulated with phorbol 12-myristate 13-acetate (PMA) and ionomycin for four hours. During the final hour of the experiment, cells were treated with 2-deoxy-D-glucose (100 mM) and oligomycin (1 μM), either separately or in combination, for approximately 45 minutes. In the final fifteen minutes of the experiment, puromycin was added at a concentration of 10 μg/ml. Following the conclusion of the treatment period, the cells were washed with cold PBS and stained using an Fc receptor blockade and Zombie Aqua as the death cell marker. Following another wash, the cells were fixed, permeabilized, and subjected to intracellular staining as previously described, incorporating an anti-puromycin monoclonal antibody conjugated with Alexa Fluor 488 (Biolegend, Cat. No.: 381506; 1:200).

### Glucose uptake assay

A total of approximately 10^6^ cells were harvested and washed with FACS buffer. Subsequently, the cell pellets were reconstituted in a standard R10 medium prepared with Glucose-free RPMI1640, and 300 μM of 6-NBDG was introduced to the samples. Following a 30-minute incubation period at 37°C, the samples were washed again with FACS buffer in preparation for flow cytometry analysis.

### Extracellular Flux Analysis

Extracellular flux analysis was performed using a Seahorse XFe96 device following the previously published protocol (Abir et al., 2024) and the Seahorse protocol provided by Agilent Inc. In summary, After PMA/ionomycin restimulation, cells were collected and washed with their corresponding Glyco-stress test (GST), Mito-stress test (MST), and the ATP analysis medium. About 1.8 X 10^5^ cells were seeded to the seahorse 96 well cell culture plate coated with Poly-D-Lysine and after 45 min of incubation in a CO2-free incubator, they were subjected to experiment. The data was analyzed with the Agilent Wave software.

### Glucose & Lactate measurement

The supernatant from cultures of naïve CD4^+^ cells, activated T cells, and Th1 cells were collected on the last day. The glucose and lactate levels in the medium were determined using the Super GL compact instrument (Hitado GmBH), as described previously (Urbanczyk et al., 2022). Before the introduction of the samples into the measuring Eppendorf, the instrument was calibrated with Super GL calibration fluid. With the help of a capillary, the test sample was introduced to the Glucocapil reaction vessels, which were then introduced to the device. Subsequently, the device automatically calculated and displayed the results.

### Immunoprecipitation (IP) and Western Blot

Approximately 10⁷ viable cells were lysed in NP40 cell lysis buffer (1% Nonidet P-40, 150 mM NaCl, 5 mM EDTA, 50 mM Tris/HCl pH7.4) containing 1 mM phenyl-methyl-sulfonyl-fluoride (PMSF) and 100 mM NaVO₃. The cells were lysed on ice for 10 minutes, followed by a 10-minute centrifugation at 10,000 x g and 4C. Supernatants were collected and incubated with 5 µl of mouse αSTAT1 and 50 µl of Protein G Sepharose for 2 hours at 4°C on a mixing rotor. The control sample was prepared by incubating 5 µl of the antibody with 50 µl Protein G Sepharose, in buffer only. Following the incubation period, the lysate was washed twice with NP40 buffer and boiled at 65°C for five minutes in 1 x SDS-sample loading buffer (2% SDS, 62.5 mM Tris/HCl pH6.8, 10% glycerol, 100 mM di-thio-threitol). Samples were subjected to 10% SDS-PAGE and western blotting to nitrocellulose for 45 minutes at 400mA and 25V. The membrane was stained with Ponceau S for 2 minutes and rinsed with Tris-buffered saline (50 mM Tris/HCl, pH7.4, 150 mM NaCl) including 0.1% Tween-20 (TBST) for 15 minutes. Then the membrane was blocked with 5% skimmed milk in TBST for an hour. The membrane was washed again for 30 minutes and stained with primary and secondary antibodies for an hour, followed by a 4 x 5 min washes with TBST. The membrane was developed with an in house enhanced chemiluminescence (ECL) kit for 2 minutes and exposed to X-ray films. Developed X-ray films were scanned, converted to TIFF and analyzed with ImageJ.

### Homology directed repair (HDR) DNA template

The HDR templates were synthesized by Twist Bioscience, CA. The left homology arm (LHA; 386 bp) is followed by the wild-type sequence of STAT1 exon 18 or an edited sequence containing the amino acid point mutations S499A and T510A. The subsequent self-cleaving peptide P2A separates the *stat1* sequence from an EGPF reporter. After the stop codon (TAA), the 376 bp right homology arm (RHA) concludes the HDR template (Figure 7). The sequences of these segments are depicted in Table S1. The DNA construct was delivered as a sequence-verified plasmid. The lyophilized plasmid was reconstituted with sterile water to 100 ng/μL and amplified by PCR to generate a linearized double-stranded HDR template. Each 50 μl PCR reaction contained 5 x Q5 Reaction Buffer, 0.5 μM STAT1 HDR genomic forward primer targeting LHA (5’-AGAGGTGAAACAGGAAGCGAG-3’), 0.5 μM STAT1 HDR genomic reverse primer targeting RHA (5’-CTTTCCCTTGGGAATTCATCTCAG-3’), 0.2 mM dNTPs, 0.5 μL Q5 DNA Polymerase, and 200 ng reconstituted DNA in PCR grade water. The PCR was run with the following cycling conditions: Initial denaturation at 98°C for 5 min, 30 cycles of 98°C for 10 sec, 60°C for 30 sec and 72°C for 3 sec, final elongation at 72°C for 2 min, and hold at 4°C. Successful amplification was confirmed with an 1% agarose gel and amplified HDR template was purified with a MinElute PCR Purification Kit (Qiagen, 28004) according to the manufacturer’s instructions.

### Ribonucleoprotein (RNP) production

In brief, 40 μM gRNAs were produced by mixing equimolar amounts of trans-activating crRNA (tracrRNA) (Integrated DNA Technologies, 1072534) with STAT1 crRNA (5’ - GACTCCACCATGTGCACGAT-3’, Integrated DNA Technologies), incubated at 95°C for 5 min with subsequent cool down to room temperature. Afterwards, 50 μg/sample poly-L-glutamic acid (PGA; Sigma-Aldrich, P4761) (1,2) and 20 μM electroporation enhancer (Integrated DNA Technologies, 10007805) were added to the gRNA. RNP production was concluded by adding equal volume of Cas9 Nuclease V3 (Integrated DNA Technologies, 1081059, diluted to 6 μM) to the gRNA (40 μM) respectively. RNPs were incubated for 15 min at RT and subsequently stored on ice for processing at the same day (Kath et al., 2022; Nguyen et al., 2020).

### Electroporation and cultivation of edited cells

Naïve CD4^+^CD45RA^+^ cells were cultured in activation medium or differentiation medium (early differentiation) for 24 hours and subjected to electroporation. Prior to electroporation, the DNA-sensing inhibitor RU.521 (InvivoGen, inh-ru521) (1) was added to the cells at a final concentration of 4.82 nM for 6 h. Afterwards, activation was stopped by transferring cells to a new plate in fresh complete RPMI medium. For electroporation (Shy et al., 2023), 1×10^6^ activated cells per electroporation sample were resuspended in 20 μL P3 electroporation buffer (Lonza, V4SP-3960) and then mixed with DNA/RNP mix (0.5 μg HDR template, 3.5 μL STAT1 RNPs). After transfer into the 16-well Nucleocuvette^TM^ Strip (Lonza, V4SP-3960), cells were electroporated (pulse sequence EH100) in the Lonza 4D-Nucleofector^TM^. After electroporation, cells were seeded in 900 μl of antibiotic-free RPMI medium supplemented with 180 U/mL IL-2. After 15 min, 20 μl of a mixture containing 0.5 μM HDAC class I/II Inhibitor Trichostatin A (AbMole, M1753) and 10 μM DNA-dependent protein kinase (DNA-PK) inhibitor M3814 (chemietek, CT-M3814) were added to each sample (3). Cells were incubated for 12-18 h in 24-well plates and then cells were cultivated in completeRPMI medium and 10ng/mL IL12. A subset of activated cells was further cultured under differentiation conditions post-electroporation, representing the late differentiation stage. GFP⁺ cells were identified as successfully edited cells. Cells edited with Ser499 and Thr510 mutated construct are marked as STAT1^mut^ and wild-type constructs are marked as STAT1^wt^. Experiments were performed with 3 healthy donor samples with 3-10 technical replicates. Flow cytometry analysis was performed to compare GFP⁺IFNγ⁺ cells with GFP⁻IFNγ⁺.

## Statistical analysis

GraphPad Prism 9 was used for the statistical analysis (GraphPad Software, San Diego, CA, USA). Two groups of data were analyzed using the Wilcoxon Matched Paired T-test. Group data was analyzed with one-way or two-way ANOVA with Turkey’s/Dunnett multiple comparisons. The graphs were either violin plots or bar diagrams with standard deviation.

## Abbreviations

AAO: amino acid oxidation
Act.T: activated T cells
2-DG: 2-Deoxyglucose
ECAR: extracellular acidification rate
FAO: fatty acid oxidation
HDR: homology directed repair
IFN: Interferon
IL: Interleukin
JAK: Janus kinase
FI: fluorescence intensity
MFI: median fluorescence
MSC: mesenchymal stem cell
N.CD4^+^: naïve CD4^+^ T cells
6-NBDG: (6-(*N*-(7-Nitrobenz-2-oxa-1,3-diazol-4-yl)amino)-6-Deoxyglucose)
OGA: O-Glycanase
OGT: O-Glycosyltransferase
OxPhos: oxidative phosphorylation
PBMC: peripheral blood mononuclear cells
PMA: Phorbol-12-myristat-13-acetat
SCENITH: single cell energetic metabolism by profiling translation inhibition
SHP2: Src homology region 2 domain-containing phosphatase-2
STAT: signal transducer and activator of transcription
Th: T helper

## Notes

### Competing Interest Statement

The authors have declared no competing interest.

