## Supplementary material for "Glycolytic flux sustains human Th1 identity and effector function via STAT1 glycosylation": Abir_2025_Supplement

**General gating strategy**

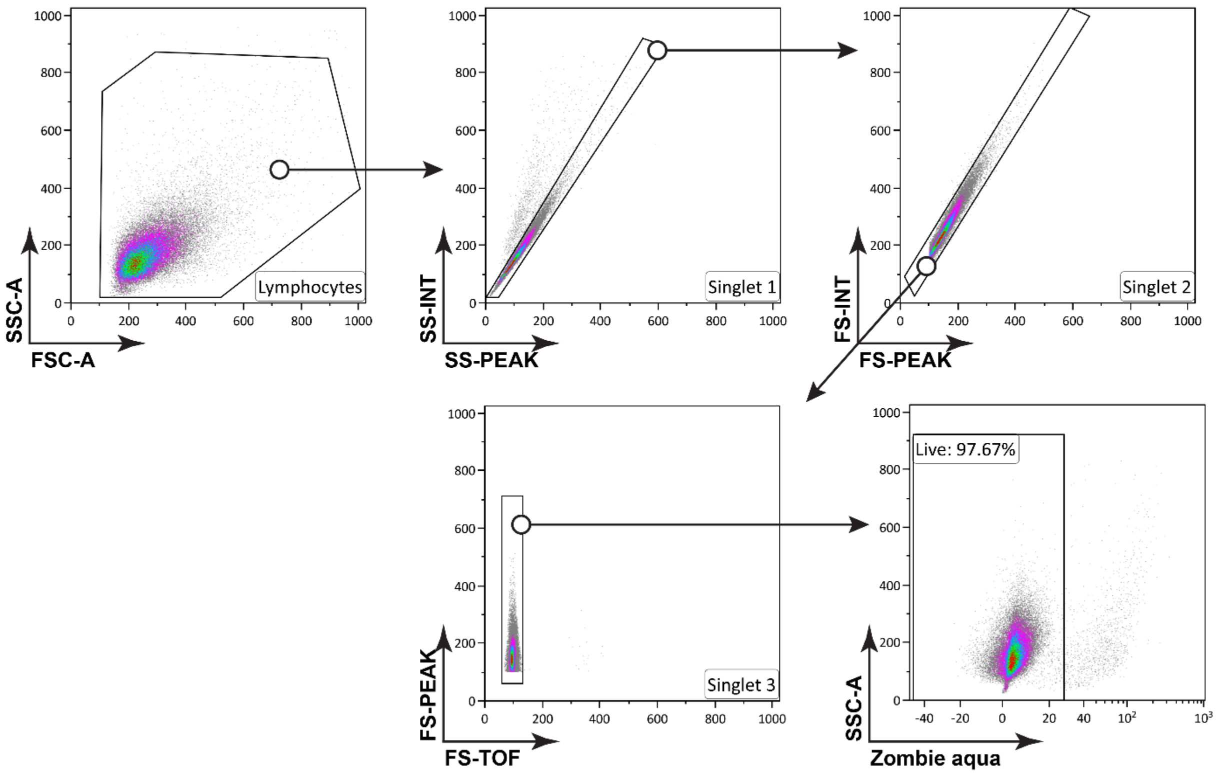


### Figure S1: General gating strategy for flow cytometric analysis

Events are plotted against the forward scatter (FSC) and the side scatter (SSC) to detect the lymphocyte population and then were gated against different combinations of forward scatter (FSC) and the side scatter (SSC) to identify singlets. Live cells were separated from dead cells by plotting against the Zombie Aqua where Zombie Aqua$^{+}$ cells were considered dead. Further analysis was performed with the Zombie Aqua$^{-}$ cells with their corresponding markers unless mentioned otherwise.

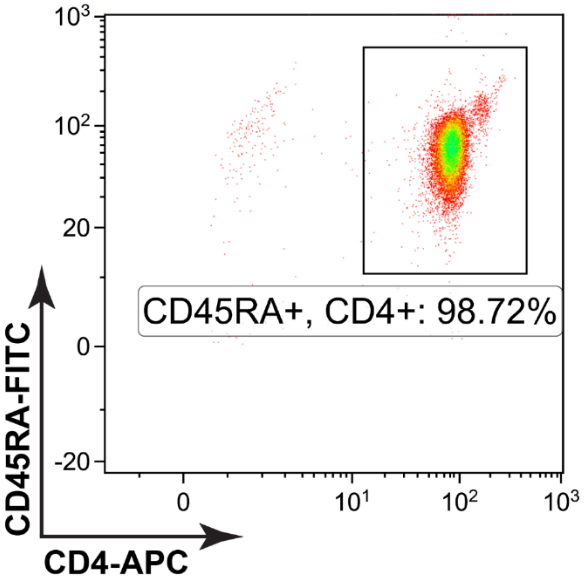


**Figure S2: Assesment of $CD4^+CD45RA^+$ purity**

Untouched naïve $CD4^+CD45RA^+$ cells were isolated from healthy donor PBMC using a MACS Naive $CD4^+$ T Cell Isolation Kit II from Miltenyi. The efficacy of the isolation was assessed by staining the cells with Anti-CD4 antibody conjugated with APC and Anti-CD45RA antibody conjugated with FITC as shown in the representative dot plot.

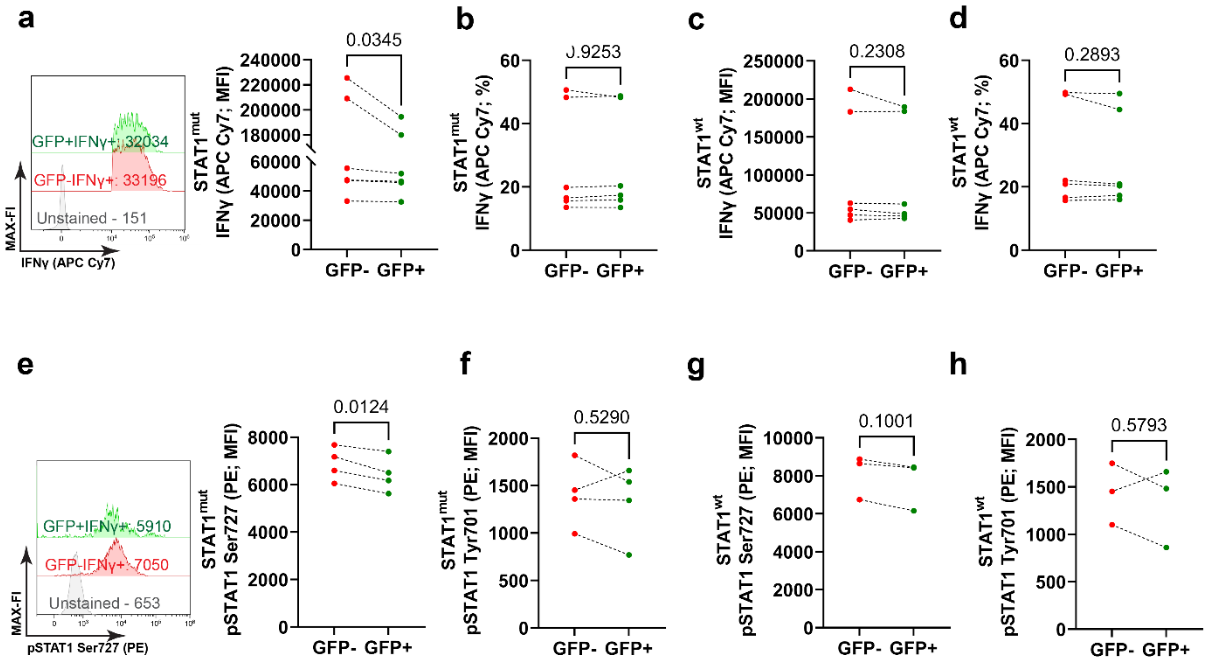


**Figure S3: Analysis of Th1 effector function and STAT1 Phosphorylation upon mutation of the STAT1 glycosylation (O-Glycosylation ) sites, late differentiation**

a. Representative histograms with MFI of the anti-IFNγ antibody conjugated with APC Cy7, GFP$^+$ or GFP$^-$ populations of STAT1$^{mut}$ cells, n=6. b. Frequencies of IFNγ producing cells gated from GFP$^+$ or GFP$^-$ populations of STAT1$^{mut}$ cells, n=6. c. MFI of GFP$^+$IFNγ$^+$ and GFP$^-$IFNγ$^+$ cells from the STAT1$^{wt}$ group, n=6. d. Frequencies of IFNγ producing cells gated from GFP$^+$ or GFP$^-$ populations of STAT1$^{wt}$ cells, n=6. e. MFI of STAT1 Ser727 (PE) of GFP$^+$IFNγ$^+$ and GFP$^-$IFNγ$^+$ cells from the STAT1$^{mut}$ group, n=4. f. MFI of STAT1 Tyr701 (PE) of GFP$^+$IFNγ$^+$ and GFP$^-$IFNγ$^+$ cells from the STAT1$^{wt}$ early differentiation group, n=5. g. MFI ofSTAT1 Ser727 (PE) of GFP$^+$IFNγ$^+$ and GFP$^-$IFNγ$^+$ cells from the STAT1$^{wt}$ group, n=3. h. MFI ofSTAT1 Tyr701 (PE) of GFP$^+$IFNγ$^+$ and GFP$^-$IFNγ$^+$ cells from the STAT1$^{wt}$ group, n=3. Data were analyzed with a paired t-test, p-values are indicated on top of each graph.

**Table S1: Sequences of the homology directed repair templates**

| **Name** | **Sequences** |
|---|---|
| **LHA** | AGAGGTGAAACAGGAAGCGAGTGTCATTTTGTTTGGCTTCAGTTGG<br>AAGCGTGTTAAGAGACTCGAATTCTTTGCTGCTGTGTGCTGCTGTGT<br>GTGCACGGGTGTGTCTTCAAATGACCCCAAAGATGCCATGTATTTAG<br>ATTTTGGAGTTACAAGTAAATTTTAGTGAGTAAACAAGTTCACAAAT<br>GTGCATCTGTAAATAATGAAAATTGACTGTATTTCTCTTCCCCTACTGT<br>GAAAGCACCTGTGTGTCATATAAACTAGAATTGAACTTTGGGATGGA<br>CATATGTTTTAGTGCCACACTTGTGACTGGTGTCTCTGTAGTAACCCT<br>TAGATTTTGGGTGTTTTCTCTCTAGAATCTGTCCTTCTTCCTGACTCCA<br>CCATGTGCA |
| **STAT1 WT insert** | AGGTGGGCTCAGCTGTCAGAAGTGCTGTCCTGGCAGTTCTCTAGCG<br>TCACCAAAAGAGGGCTCAATGTTGACCAACTGAACATGTTGGGCGA<br>GAAGCTTTTAGGA |
| **STAT1 edited insert** | AGGTGGGCTCAGCTGGCCGAAGTGCTGTCCTGGCAGTTCTCTAGCG<br>TCGCAAAAAGAGGGCTCAATGTTGACCAACTGAACATGTTGGGCGA<br>GAAGCTTTTAGGA |
| **P2A** | GGCAGCGGCGCCACCAACTTCAGCCTGCTGAAGCAGGCCGGCGAC<br>GTGGAAGAGAACCCCGGGCCC |
| **EGFP** | ATGGTGAGCAAGGGCGAGGAGCTGTTCACCGGGGTGGTGCCCATC<br>CTGGTCGAGCTGGACGGCGACGTAAACGGCCACAAGTTCAGCGTG<br>TCCGGCGAGGGCGAGGGCGATGCCACCTACGGCAAGCTGACCCTG<br>AAGTTCATCTGCACCACCGGCAAGCTGCCCGTGCCCTGGCCCACCC<br>TCGTGACCACCCTGACCTACGGCGTGCAGTGCTTCAGCCGCTACCCC<br>GACCACATGAAGCAGCACGACTTCTTCAAGTCCGCCATGCCCGAAGG<br>CTACGTCCAGGAGCGCACCATCTTCTTCAAGGACGACGGCAACTACA<br>AGACCCGCGCCGAGGTGAAGTTCGAGGGCGACACCCTGGTGAACCG<br>CATCGAGCTGAAGGGCATCGACTTCAAGGAGGACGGCAACATCCTGG<br>GGCACAAGCTGGAGTACAACTACAACAGCCACAACGTCTATATCATGG<br>CCGACAAGCAGAAGAACGGCATCAAGGTGAACTTCAAGATCCGCCAC<br>AACATCGAGGACGGCAGCGTGCAGCTCGCCGACCACTACCAGCAGAA<br>CACCCCCATCGGCGACGGCCCCGTGCTGCTGCCCGACAACCACTACCT<br>GAGCACCCAGTCCGCCCTGAGCAAAGACCCCAACGAGAAGCGCGATC<br>ACATGGTCCTGCTGGAGTTCGTGACCGCCGCCGGGATCACTCTCGGCA<br>TGGACGAGCTGTACAAGTAA |
| **RHA** | CGATGGGCTCAGCTTTCAGAAGTGCTGAGTTGGCAGTTTTCTTCTGTCA<br>CCAAAAGAGGTCTCAATGTGGACCAGCTGAACATGTTGGGAGAGAAGC<br>TTCTTGGTAGAGAAGCTTCTTGGTATATGCATATTAACTTGTTATGTTTATA<br>AAAATTGAAATTCATAAAAATATCTCTCTAATTGCTCTTTTCCCCTCTGCTA<br>TTTTGTTAAAGGTAAAAAAGTACTAAAATCTGTCAGCTTTTCAAGCTATAG<br>TTTATTATAGCTAAGTGAGAATCATATGTCACCTTAGAAAGAAATATAGACC<br>TGATAACATTTAAATGAATCCGTTCCTCATTGTCTCATATTAAGATTTCTGA<br>GATGAATTCCCAAGGGAAAG |

**Supplemental Material**

| Products | Manufacturers | Catalogue No. |
|---|---|---|
| 2-Desoxy-D-Glucose (2DG). | Sigma - Aldrich | D8375 |
| 2-Mercaptoethanol | Gibco™, Thermo Fisher Scientific | 21985023 |
| 3PO | Sigma - Aldrich | SML1343-5MG |
| Agarose | PeqLab | 732-2789 |
| Albumin fraction V ( BSA ) | Carl Roth | 8076.4 |
| Albutein 50 g/l Infusionslösung | Grifols Deutschland GmbH | 10408446 |
| Antimycin A | Sigma - Aldrich | A - 8674 |
| APS | Carl Roth | 9592.3 |
| BD Cytofix™ Fixation Buffer | BD | 554655 |
| BD GolgiPlug™ | BD | 555029 |
| BD Phosflow™ Perm Buffer III | BD | 558050 |
| BSA, fatty acid-free | Sigma - Aldrich | A3803 |
| DAPI | Carl Roth | 28717-90-3 |
| DMSO | Molecular Probes | C10634 |
| Ethidium bromide | Carl Roth | 2218 |
| EtOH | Merck | 1.00983.2500 |
| FCCP | Sigma - Aldrich | C2920 |
| Fetal Bovine Serum | Gibco™, Thermo Fisher Scientific | 16000044 |
| Glucose | Carl Roth | HN06.3 |
| Glutaraldehyde | Carl Roth | 4995.1 |
| Glycerin | ROTIPURAN®, Carl Roth | 200-289-5 |
| Hydrochloric acid fuming 37% | Merck Millipore | 113386 |
| Ionomycin calcium salt | Sigma - Aldrich | I3909-1ML |
| L-Glutamine (200 mM) | Gibco™, Thermo Fisher Scientific | 25030081 |
| Milk Powder | Carl Roth | T145.3 |

| | | |
|---|---|---|
| Mowiol 4-88 | Calbiochem | 47-590-4100GM |
| $NaN_3$ | Carl Roth | 26628-22-8 |
| NP 24 - KLH | Biosearch Technologies | N - 5060-5 |
| Oligomycin | Sigma - Aldrich | 75351 |
| OSMI1 | Sigma - Aldrich | SML1621-5MG |
| PageRuler Prestained | Thermo Scientific | 26616 |
| p-Cumaric acid | Carl Roth | 9908.2 |
| Penicillin - streptomycin | Gibco | 15140-122 |
| Penicillin-Streptomycin (10,000 U/mL) | Gibco™, Thermo Fisher Scientific | 15140122 |
| Phosphate - Buffered Saline (PBS), 1x | Gibco™, Thermo Fisher Scientific | 14190-094 |
| PMSF | Sigma - Aldrich | 329-98-6 |
| Poly-D-Lysine | Sigma - Aldrich | P6407 |
| Ponceau S | Carl Roth | 5938.1 |
| Proteinase K | Peqlab | 04-1076 |
| Rotenone | Sigma - Aldrich | R8875 |
| RPMI1640 | Gibco™, Thermo Fisher Scientific | 11875093 |
| SDS ultra pure | Carl Roth | 205-788-1 |
| Seahorse XF RPMI medium, pH 7.4 | Agilent Technologies | 103576-100 |
| Sodium chloride | Carl Roth | 231-598-3 |
| Sodium hydroxide | Merck Millipore | 106462 |
| Sodium Pyruvate | Gibco™, Thermo Fisher Scientific | 11360070 |
| TEMED | Carl Roth | 2367.3 |
| Tris | Carl Roth | 4855.3 |
| Trizma Base | Sigma-Aldrich | T1503 |
| Tween - 20 | Carl Roth | P1379 |

**Table 2: List of commercial kits**

| Products | Manufacturers | Catalog No. |
|---|---|---|
| Naive CD4+ T Cell Isolation Kit II, human | Miltenyi | 130-094-131 |
| PyroMAT™ System | Merck Millipore | D5047 |
| Seahorse Extracellular Flux Analysis Kit | Agilent Technologies | 103793-100 |
| Seahorse XF Real-Time ATP Rate Assay Kit | Agilent Technologies | 103592-100 |
| VersaComp Antibody Capture Kit | Beckman Colter Life sci | B22804 |

**Table 3: PCR**

| Products | Manufacturers | Catalog No. |
|---|---|---|
| DNA/RNA dye, peqGREEN | VWR Peqlab | 732-3196 |
| dNTP Mix (10 mM each) | Thermo Fisher Scientific | R01922 |
| GeneRuler 1 kb DNA Ladder | Thermo Fisher Scientific | SM0311 |
| Q5 high-fidelity polymerase | New England Biolabs | M0491S |
| MinElute PCR Purification Kit | Qiagen | 28004 |
| RT-PCR Grade Water | Thermo Fisher Scientific | AM9935 |

**Table 4: List of antibodies and cytokines**

| Products | Manufacturers | Catalog No. |
|---|---|---|
| Alexa Fluor 647 Anti-O-Linked N-Acetylglucosamine antibody [RL2] (ab201994) | Abcam | ab201994 |
| Alexa Fluor 647 Mouse anti-Total Stat1 (N-Terminus) | BD Biosciences | 558560 |
| APC anti-human CD4 Antibody | BioLegend | 317416 |
| APC/Cyanine7 anti-human IFN-γ Antibody | Biolegend | 506524 |
| BD Pharmingen™ Human BD Fc Block™ | BD | 564219 |
| Brilliant Violet 421™ anti-human IL-4 Antibody | BioLegend | 500826 |
| FITC anti-human CD3 Antibody | BioLegend | 300406 |
| FITC anti-human CD45RA Antibody | BioLegend | 304148 |
| FITC anti-human IL-17A Antibody | BioLegend | 512304 |
| FITC OGT1 Polyclonal Antibody | Invitrogen | OGT1-FITC |
| FITC Phospho-Jak2 (Tyr1007, Tyr1008) Recombinant Rabbit Monoclonal Antibody (JAK2Y10071008-PB6), | Thermo Fisher Sci. | MA5-37198 |
| Human IL-12, premium grade | Miltenyi Biotec | 130-096-705 |
| Human IL-2 IS, research grade | Miltenyi Biotec | 130-097-743 |

| | | |
|---|---|---|
| Human IL-4 Antibody | R&D System | MAB204-100 |
| Naive CD4+ T Cell Isolation Kit II, human | Miltenyi | 130-094-131 |
| O-GlcNAc Antibody (RL2) [HRP] | Novus Biologicals | NB300-524H |
| O-GlcNAc Antibody (RL2) [HRP] | Novus Biologicals | NB300-524H |
| O-GlcNAc Monoclonal Antibody (RL2), Alexa Fluor™ 488, eBioscience™ | Thermo Fisher | 53-9793-42 |
| PE anti-STAT1 Phospho (Ser727) Antibody | Biolegend | 686404 |
| PE Mouse Anti-Stat1 (pY701) | BD Biosciences | 612564 |
| PE Phospho-Jak1 (Tyr1022, Tyr1023) Recombinant Rabbit Monoclonal Antibody | Invitrogen | MA5-36891 |
| PerCP/Cyanine5.5 anti-T-bet Antibody | BioLegend | 644806 |
| Phospho-Stat1 (Ser727) Antibody #9177 | Cell Signaling Technology | 9177S |
| Phospho-Stat1 (Tyr701) (58D6) Rabbit mAb #9167 | Cell signaling | 9167S |
| Purified anti-human IL-4 Antibody | BioLegend | 500802 |
| Stat1 (9H2) Mouse mAb #9176 | Cell signaling | 9176S |
| Stat1 Antibody #9172 | Cell signaling | 9172S |
| STAT4 pY693 Antibody, anti-human, FITC, REAfinity™ | Miltenyi Biotec | 130-114-491 |
| Ultra-LEAF™ Purified anti-human CD28 Antibody | BioLegend | 302934 |
| Ultra-LEAF™ Purified anti-human CD3 Antibody | BioLegend | 300438 |
| Zombie Aqua™ Fixable Viability Kit | BioLegend | 423101 |

**Table 4: Buffer solutions and medium**

| **Reagent** | **Composition** |
|---|---|
| Anode buffer | $dH_2O$<br>25mM Tris<br>20ml/100ml MeOH |
| Antibody dilution | PBS |

| | |
|---|---|
| | 3g/100ml BSA<br>0.01ml/100ml Tween-20<br>0.1ml/100ml $NaCl_3$ |
| Cathode buffer | $dH_2O$<br>40mM 6-Aminocaproic acid<br>20ml/100ml MeOH |
| Cell sorting buffer (MACS) | PBS<br>2% Human Serum Albumin<br>5mM EDTA |
| ECL | 2 ml 1M Tris pH 8.5<br>17.7 ml $H_2O$<br>90 µl 90 mM p-Cumaric acid<br>200 µl 200 mM Luminol<br>6 µl $H_2O_2$ |
| FACS buffer | PBS<br>2% FCS<br>5ml 100mM EDTA |
| Ponceau S | $dH_2O$<br>1g/l Ponceau S |
| R10 Medium | RPMI1640<br>2mM L-Glutamine<br>100U/ml Penicillin<br>100U/ml Streptomycin<br>50µM 2-Mercaptoethanol |
| Seahorse ATP analysis medium | 1mM Sodium pyruvate<br>2mM L-Glutamine<br>10mM Glucose |
| Seahorse cell culture plate coating buffer | $dH_2O$<br>100ng/ml Poly-D-lysine |
| Seahorse glyco-stress test medium | 1mM Sodium pyruvate<br>2mM L-Glutamine |

| | |
|---|---|
| Seahorse mito-stress test medium | 1mM Sodium pyruvate<br>2mM L-Glutamine<br>10mM Glucose |
| TBS-T | $dH_2O$<br>40mM Tris/HCL pH 8.5<br>20mM Sodium acetate<br>1mM EDTA |

**Table 5: List of software**

| **Name - Version** | **Utilization** |
|---|---|
| Adobe Illustrator 2017 | Illustrations |
| GraphPad Prism - 9.0.0 | Statistical analysis and data presentation |
| ImageJ - Fiji - 1.53t | Western Blot analysis |
| Kaluza Analysis - 2.1 | Flow Cytometry Analysis |
| Microsoft Excel 2019 | Data management |
| Microsoft Word 2019 | Manuscript writing and presentation |
| Wave 2.6.3.5 | Seahorse analysis |
| Zotero - 6.0.30 | Citation management |
